# *De novo* clustering of long-read amplicons improves phylogenetic insight into microbiome data

**DOI:** 10.1101/2023.11.26.568539

**Authors:** Yan Hui, Dennis Sandris Nielsen, Lukasz Krych

## Abstract

Long-read amplicon profiling through read classification limits phylogenetic analysis of amplicons while community analysis of multicopy genes, relying on unique molecular identifier (UMI) corrections, often demands deep sequencing. To address this, we present a long amplicon consensus analysis (LACA) workflow employing multiple *de novo* clustering approaches based on sequence dissimilarity. LACA controls the average error rate of corrected sequences below 1% for the Oxford Nanopore Technologies (ONT) R9.4.1 and ONT R10.3 data, 0.2% for ONT R10.4.1, and 0.1% for high-accuracy ONT Duplex and Pacific Biosciences (PacBio) circular consensus sequencing (CCS) data in both simulated 16S rRNA and real 16-23S rRNA amplicon datasets. In high-accuracy PacBio CCS data, the clustering-based correction matched UMI correction, while outperforming 4×UMI correction in noisy ONT R10.3 and R9.4.1 data. Notably, LACA preserved phylogenetic fidelity in long operational taxonomic units and enhanced microbiome-wide phenotype characterization for synthetic mock communities and human vaginal samples.

## Main

High-throughput amplicon sequencing remains a cost-effective strategy to analyze genetic polymorphisms in DNA specimens, also in situations where the quantity of the sample DNA is low^1^, rendering it a widely employed technique in viral^2^ and microbial^3^ profiling. Until now, short-read technologies like Illumina dominated amplicon sequencing. However, the maximum insert size of ∼ 500 bp^4^ necessitates either fragmenting amplicons or limiting the length of an amplified region. The emergence of long-read sequencing technologies, such as Oxford Nanopore Technologies (ONT) and Pacific Biosciences (PacBio), has revolutionized amplicon sequencing, enabling the complete sequencing of amplicons up to 10 kb^5^. However, these technologies exhibit variable sequencing error rates, ranging from 1% to 15%, depending on the used platforms^6–8^. To address this, many long-read amplicon analysis approaches often resort to closed-reference classification. For instance, the official ONT EPI2ME analysis toolbox utilizes this strategy, employing read-by-read classification against a known reference database via Centrifuge^9^. More recently, Emu^10^ introduced an expectation-maximization algorithm to achieve species-level microbial profiling through full-length 16S rRNA gene amplicon sequencing. Nevertheless, these methods come with limitations, potentially overlooking species not included in trusted reference databases and failing to retain the evolutionary links of taxonomic traits in downstream analysis.

The variable sequencing error rate challenges *de novo* clustering of long noisy reads into operational taxonomic units (OTUs). IsONclust^11^ is an innovative method that leverages base quality values in *de novo* clustering of such reads. Based on this, NGSpeciesID achieved remarkable accuracy, surpassing 99.3% when compared to Sanger sequencing, in retrieving ribosomal operon sequences from distant fungal strains^12^. Using an assembly-free strategy of read binning with 5-mer frequency similarity^13^, NanoCLUST pioneered a reference-free workflow for OTU clustering of noisy ONT amplicons of full-length 16S rRNA gene^14^. NanoCLUST attained species-level resolution and effectively profiled an eight-strain mock community^14^. In high-accuracy PacBio circular consensus sequencing (CCS) data, IsoCon demonstrated nucleotide-level precision in deciphering highly similar multigene families^15^. Notably, with noisy ONT R9.4.1 transcriptome data, isONcorrect obtained a comparable median accuracy of 98.9-99.6% to PacBio CCS without reference reliance^16^. This suggests the potential to expand the use of IsoCon on noisy ONT data. Besides, the implementation of unique molecular identifiers (UMIs) in library preparation has been well-documented as an effective means to reduce the error rate in ONT and PacBio CCS sequencing to below 0.01%, but unfortunately the UMI-approach also requires rather deep sequencing depth^5^.

Until now integrated frameworks such as QIIME 2^17^ offer limited support for long-read amplicon analysis. Further, a systemic assessment of the precision and recall of these reference-free methods is lacking. To bridge this gap, this study presents a scalable and reproducible analysis workflow for *de novo* long amplicon consensus analysis (LACA). LACA incorporates multiple sequence dissimilarity clustering approaches to identify true amplification templates amidst the noise. Utilizing *in silico* benchmarking, public UMI-tagged sequencing data, and full-length 16S rRNA gene amplicon ONT sequencing data of synthetic mock and human vaginal microbiomes, our study offers a comprehensive overview of the reference-free solutions in long-read amplicon analysis.

## Results

### A reproducible and scalable workflow for long amplicon consensus analysis

LACA is designed for *de novo* OTU picking from long noisy amplicon sequencing data, regardless of whether UMIs are included in library preparation (Fig. 1a). It is modularized with Snakemake^18^ sub-workflows and offers user-friendly controls for demultiplexing, quality control, OTU picking, and downstream community analysis, i.e., quantification, taxonomy assignments, and phylogenetic reconstruction (Fig. 1b, 1c). Taking advantage of Snakemake^18^, LACA assures reproducible analysis with a pre-defined configuration file and is scalable to support large-scale sample processing in cluster systems. Unlike short-read technologies, long-read sequencing preserves the integrity of amplicons. This enables LACA to identify the amplified regions by searching for the linked primer patterns and re-orient the sequence if a reverse strand is sequenced (Fig. 1b). The remaining reads are pooled or processed independently for clustering and consensus calling after chimera and quality filtering. To avoid the heavy computing cost of pairwise global alignments, LACA borrows multiple alignment-free approaches, e.g., HDBSCAN clustering with k-mer frequency^14,13^ or error-aware clustering with minizers^11^, and consolidates the clusters with Meshclust^19^. Relying on the mean shift algorithm^20^ and alignment-free identity scores^21^, LACA identifies the centroids and members within the cluster, thus generating the consensus sequences (“kmerCon”). As shown in Fig. 3a, LACA supports refining these clusters according to alignment overlap^13^ (“miniCon”), identifying highly similar haplotypes in the cluster with isONcorrect^16^ and IsoCon^15^ (“isoCon”) and also independent molecule-level profiling with UMIs^5^ (“umiCon”). According to user requirements, the centroid is picked and polished with Racon^22^ or Medaka with the supporting reads either in a kmer-based, alignment-based, or UMI cluster. The corrected sequences are then dereplicated by sequence identity to extract representative OTUs, enabling phylogenetic inference and taxonomic assignment in community analysis (Fig. 1c). The OTU count matrix can be created by the uniquely assigned FASTQ sequence identifiers in the clustering and demultiplexing records or re-mapping the sequencing reads against the generated OTU sequences. Multiple sequencing runs of LACA outputs can be merged for meta-analysis, in which run-specific OTUs are concatenated, re-dereplicated by sequence identity, and the OTU count matrices are merged accordingly.

**Fig. 1:**
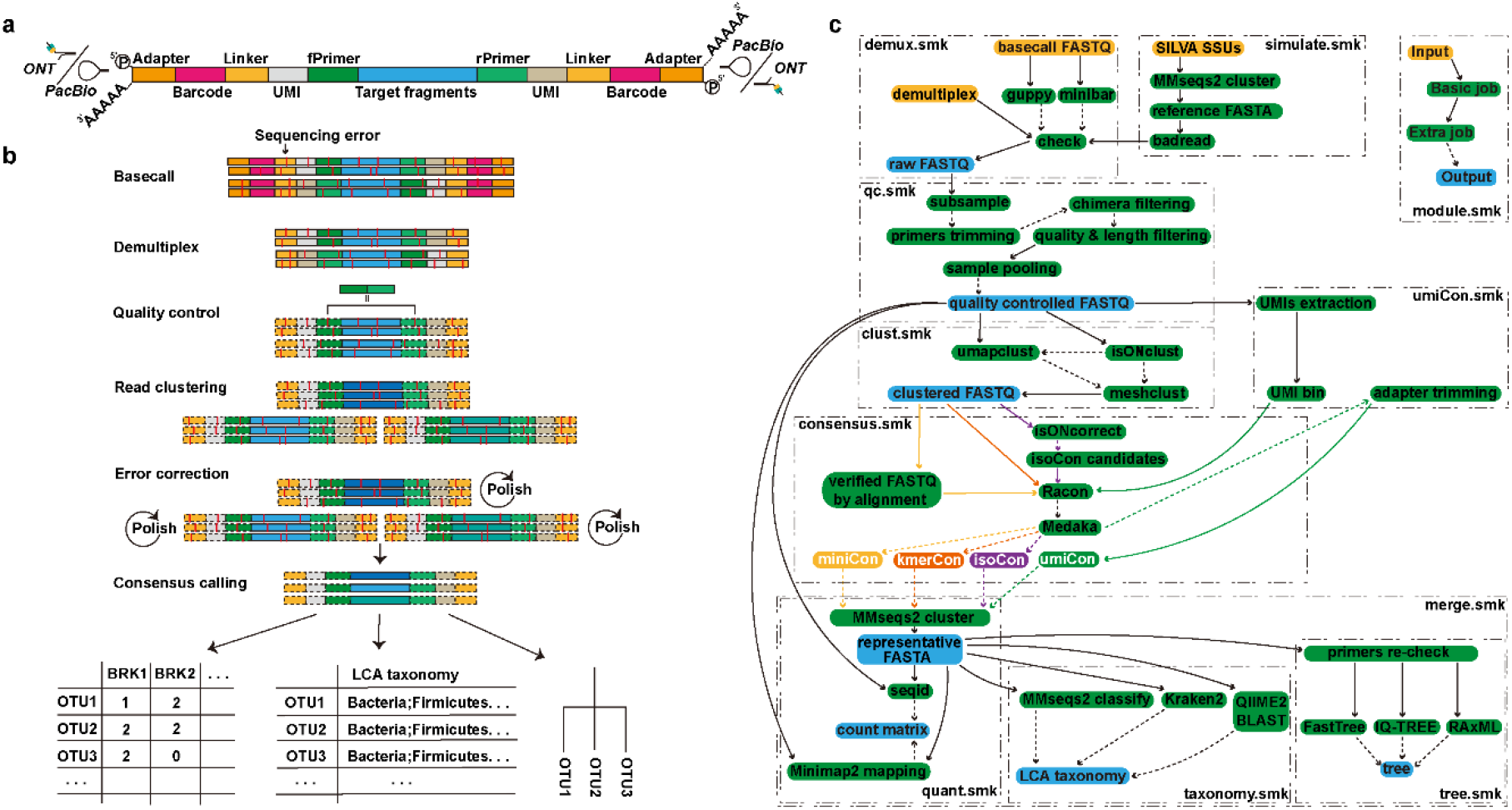
a, A schematic illustration of the amplicon structure suitable for long amplicon consensus analysis (LACA). b, c, Graphical representation of the essential steps in the LACA workflow (b) and a bioinformatic flowchart (c). Briefly, LACA enables demultiplexing sequencing reads by Oxford Nanopore Technologies with custom barcodes or directly picks external demultiplexed data for downstream analysis. The demultiplexed reads passing through primer trimming and quality control are either pooled together or processed independently to retrieve consensus sequences for OTUs picking. The OTUs are used to obtain phylogenetic and taxonomic relationships for community analysis. The OTU count table is generated by incorporating OTU clustering and read demultiplexing information, utilizing either the unique sequence ID (in a clustering-based approach) or read remapping (in a mapping-based approach). The OTUs are re-clustered to select final representatives when merging LACA runs, and the OTU table, taxonomy, and tree file are updated accordingly.

### *In silico* benchmark with SILVA amplicons

*De novo* OTU picking relies on sequence dissimilarity information to cluster amplicons from the same template. However, erroneous base calls can lead to the misclassification of similar sequences. Given the varied sequencing accuracy among long-read platforms, we posited that clustering performance on long amplicons is influenced by sequencing depth and target sequence divergence. To test this hypothesis, we selected five sets of ten small subunit (SSU) rRNA gene sequences from the SILVA database. These sequence sets had sequence divergences of 5-10%, 3-5%, 2-3%, 1-2%, and 0-1%. They served as reference sequences to simulate long amplicons from the early ONT R9.4.1 (with a mean read identity of 87.5%), ONT R10.4.1 (with a mean read identity of 95%), ONT Duplex (with a mean read identity of 99%), and PacBio CCS platforms (with a mean read identity of 99%). We initially assessed the clustering quality of alignment-free approaches on the simulated reads at an adequate sequencing coverage of 200×. We recorded the number of clusters and subsequently calculated purity and normalized mutual information (NMI) scores using the read simulation source for external cluster validation. As expected, clustering performance improved with sequencing accuracy and target sequence divergence, although various approaches exhibited different clustering sensitivity and specificity. Across all data types and simulation profiles, UMAPclust consistently generated clusters with the highest levels of purity and NMI scores compared to other methods, namely isONclust and Meshclust (Fig. 2). UMAPclust retained over 75% of the original sequencing reads except for an approximate 20% decline in identifying ONT2020 reads from highly similar reference sequences with sequence divergence below 1% (Supplementary Fig. 1 and 2). However, due to the combination of forward and reverse complement motifs in the 5-mer frequency calculation, UMAPclust lost its ability to identify sequence notation, resulting in clusters of around ten sequences rather than twenty (Supplementary Fig. 3). When combined with isONclust or Meshclust, UMAPclust split sequences based on strand orientation, doubling the clustering purity (Fig. 2). For high-accuracy ONT Duplex and PacBio CCS reads, the combination of isONclust and UMAPclust successfully generated 20 clusters with NMI scores equal to 1 in both even and skewed simulation scenarios when the target sequence divergence was above 1%. However, when clustering reads from highly similar targets (sequence divergence < 1%), the cluster purity decreased by 0.25 and 0.5 according to target sequences in cases of even or skewed abundance. Using Meshclust to consolidate UMAPclust clusters exhibited comparable NMI scores in the two high-accuracy datasets, while this combination outperformed others in clustering noisy ONT reads. Following the identity scores estimated by Meshclust, it precisely clustered ONT2020 and ONT2023 reads from target sequences with sequence divergences above 5% and 2%, and it maintained clustering purity above 0.75 when the target sequence divergence decreased to 3% and 1%, respectively. All methods demonstrated a noticeable decline in purity in the clustered ONT2023 reads with a target sequence divergence below 1%, while this value increased to 2% for the simulated ONT2020 reads. The sequential use of isONclust, UMAPclust, and Meshclust did not result in a significant improvement in cluster purity; instead, it fragmented the true clusters, though without severely hampering the NMI score. The selection of identity scores influenced both clustering accuracy and the yield of Meshclust. When employing the estimated identity score derived from the dataset, Meshclust retained approximately 75% of all types of long reads in the test. Clustering purity improved with the defined Meshclust identity score (Supplementary Fig. 4), while a corresponding reduction in the yield of retained reads was observed (Supplementary Fig. 5).

**Fig. 2.**
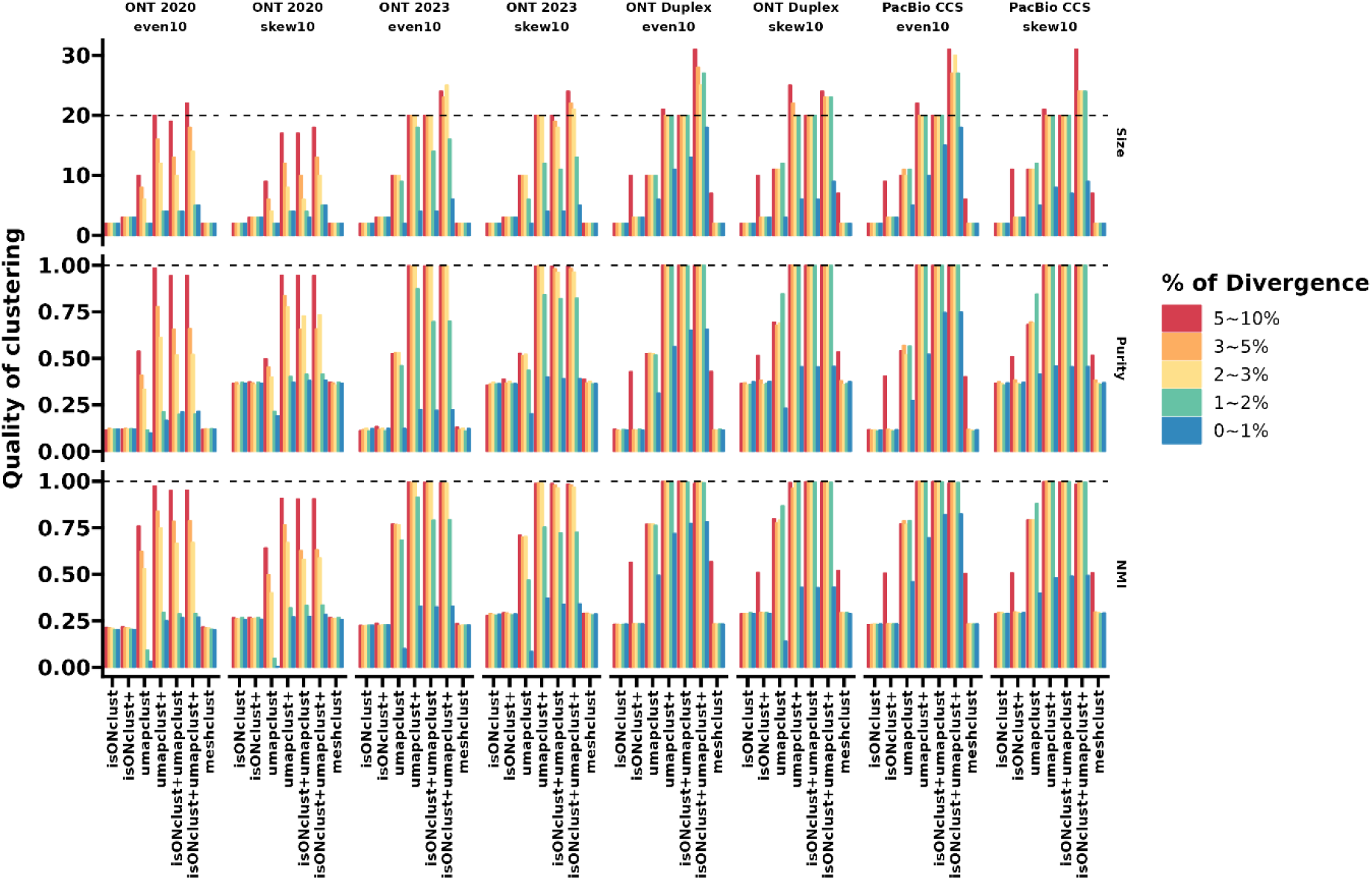
The clustering quality of various alignment-free approaches on the SILVA amplicons under the *in silico* sequencing depth of 200×. The clustering quality is assessed by size (the number of clusters), purity, and normalized mutual information (NMI). The black dashed lines suggest the theoretically optimal values, and the sequence orientation (strand) information is considered in the clustering evaluation, resulting in the theoretical number of reference sequences of 20. The simulation template references are chosen from SILVA small subunits sequences with the divergence of 5-10%, 3-5%, 2-3% 1-2%, and 0-1%, and respective results are depicted by color. Even10 refers to a set of ten reference sequences with equal abundance; skew10 refers to ten reference sequences with two sequences 10 times as abundant as the rest. The plus sign suggests the clusters are validated with Meshclust.

We benchmarked consensus calling of kmerCon, miniCon, and isoCon across a range of sequencing depths, spanning from 40× to 400× coverage. Except minCon, which exhibited a high rejection rate with noisy ONT2020 reads, LACA utilized over 75% of the reads for consensus calling, rarely incorporating chimeric sequences (Supplementary Fig. 6). And the sequencing identity of retained reads closely followed the defined distribution (Supplementary Fig. 7). Recall of target sequences generally correlated with the clustering ability of various approaches. However, when the composition of target sequences was skewed, it resulted in a retardant recall of the complete set (Fig. 3). In the kmerCon mode, consensus sequences were generated based on the alignment-free read clusters. With 100× coverage, this approach could distinguish target sequences with divergences greater than 5%, 3%, 1%, and 1% for ONT2020, ONT2023, ONT Duplex, and PacBio CCS data, respectively (Figure. 3a). Clustering purity and recall improved with sequencing depth as long as the target sequence divergence remained within the discernible range. The miniCon mode generated consensus sequences using refined clusters through alignment overlap. However, we did not observe significant improvements relative to the kmerCon mode. The isoCon mode proved most effective in retrieving the target sequence set. When sequencing depth exceeded 80× coverage, it successfully recovered all ten highly similar sequences (with sequence divergence less than 1%) in all “even10” simulated datasets except the noisy ONT2020 dataset (Fig. 3b and Supplementary Fig. 9). However, the clustering process might fragment the true clusters, and the number of clusters steadily increased with sequencing depth, exceeding 20 clusters at 400× coverage. This resulted in an approximate 10-30% decline in NMI scores across various benchmark profiles and data types (Supplementary Fig. 8). The re-clustered reads by isoCon demonstrated strong concordance with the simulation source, with cluster purity plateauing at 1 for any type of simulated ONT Duplex and PacBio CCS data and showing a slight drop in the ONT2023 data (Fig. 3a). In the ONT 2020 dataset, isoCon exhibited a cluster purity above 0.85 when identifying simulated reads from target sequences with sequence divergence above 1% at 400× coverage.

**Fig. 3:**
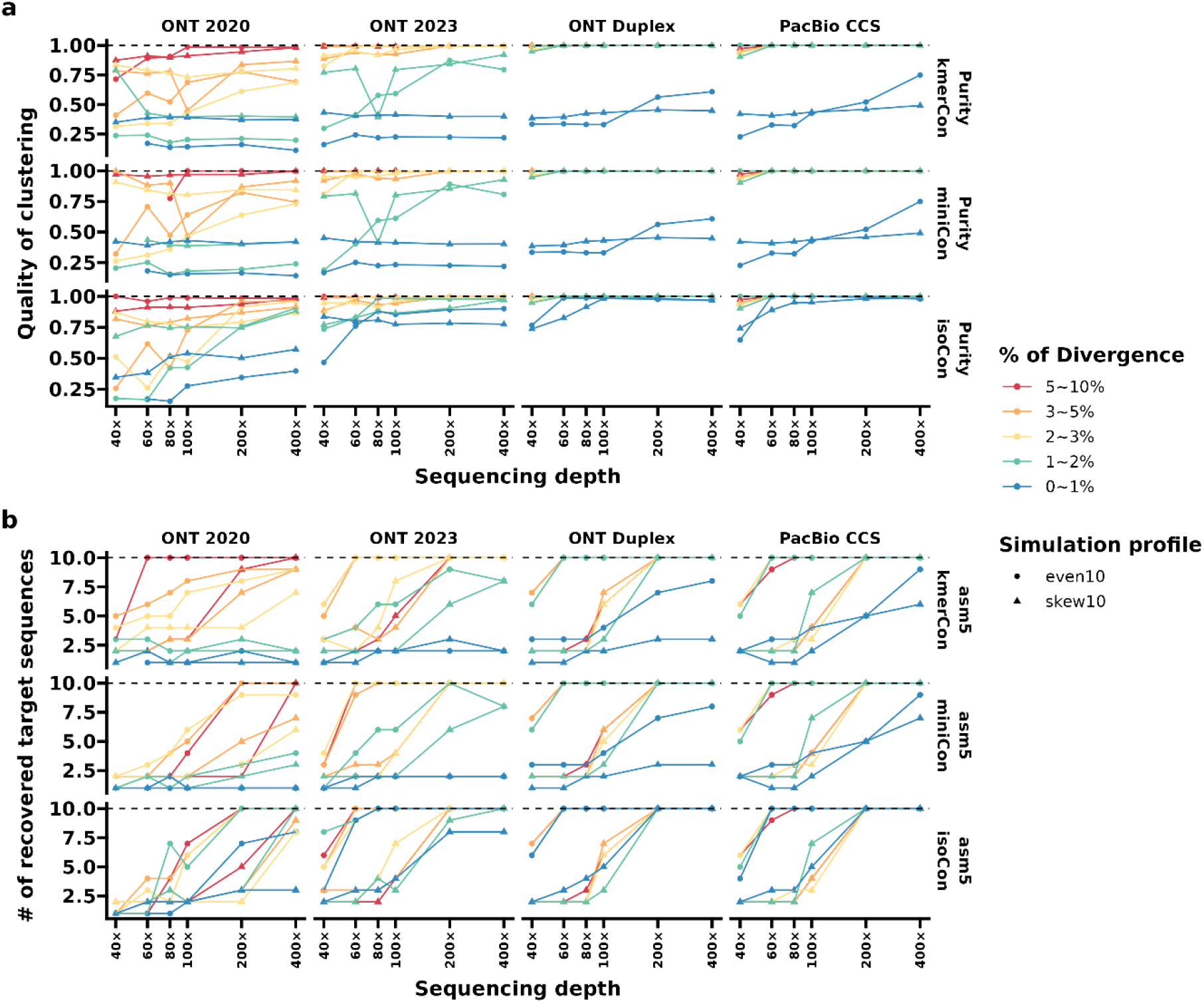
The clustering purity (a) and the number of recovered target sequences (b) of the three consensus calling approaches on the SILVA amplicons under the *in silico* sequencing depth from 40× to 400×. The black dashed lines suggest the theoretically optimal values, and the sequence orientation (strand) information is considered in the clustering evaluation, resulting in the theoretical number of reference sequences of 20. The simulation template references are chosen from SILVA small subunits sequences with the divergence of 5-10%, 3-5%, 2-3% 1-2%, and 0-1%, and respective results are depicted by color. The point shapes indicate the simulation scenarios of reference sequences in even (dots) and skewed (triangles) composition. KmerCon, consensus calling on clustered sequences with UMAPclust and Meshclust; miniCon, consensus calling on clustered sequences refined by overlap check; isoCon, consensus calling of detected isoforms by IsoCon on the clustered sequences. Even10 refers to a set of ten reference sequences with equal abundance; skew10 refers to ten reference sequences with two sequences 10 times as abundant as the rest.

The quality of corrected sequences was primarily influenced by sequencing accuracy rather than sequencing depth. All corrected sequences exhibited significant fidelity to the reference sequence, showing a minimal alignment identity of 99% and an unalignment rate of 2% in most corrected ONT2020 sequences, and 99.9% and 1% for those derived from other datasets, respectively (Supplementary Figs. 10, 11, and 12). At 200× coverage, LACA controlled the average error rate (sum of variation rate in alignments and unalignment rate) of corrected sequences to 1%, 0.2% and 0.05% for ONT R9.4.1, ONT R10.4.1 and high accuracy ONT and PacBio data, respectively (Supplementary Table 1). Notably, due to the utilization of Medaka, consensus-based corrections demonstrated the most robust performance in ONT Duplex data. The alignment error rate and the rate of unaligned regions in corrected and reference sequences were consistently below 0.1% (Supplementary Fig. 11, 12, and 13). Most corrected sequences covered over 99% of the reference sequences, except for a slight decline observed in the corrected ONT2020 sequences, typically ranging from 95% (Supplementary Fig. 13). In our alignment comparisons, we observed that the overlap-refining strategy employed by miniCon improved the quality of consensus sequences, particularly in the case of noisy ONT2020 data. However, its impact diminished as sequencing accuracy improved. The implementation of Meshclust consistently ensured the quality of consensus calling. All corrected sequences obtained from ONT Duplex and PacBio CCS data demonstrated remarkably high alignment identity, surpassing 99% when compared to the source sequences (Supplementary Fig. 11 and Supplementary Table 2). Under a more lenient control of Meshclust, supervised clusters by isONclust and UMAPclust led to a decline in identity between consensus sequences and source sequences at shallow depth, even when high-accuracy ONT Duplex and PacBio CCS reads were used (Supplementary Table 3).

### Error profiling of clustering-based and UMI-binned consensus sequences

We extended our support for consensus calling by incorporating user-defined UMI patterns and compared these *de novo* clustering approaches with molecule-level UMI corrections in a public long-read UMI amplicon dataset of 16S-23S rRNA operons sequenced via PacBio CCS, ONT R9.4.1, and R10.3 platforms. We subsampled 100,000 and 1,000,000 reads for each type of long-read amplicon data. Compared to our *in silico* tests, the alignment-free clusters were further refined into smaller micro-clusters by isoCon in the longer amplicon dataset (∼4300 bp) (Fig. 4a). In the case of shallow sequencing with 100,000 reads, the 16×isoCon sequences exhibited equivalent or even superior recall of mock operons compared to 4×UMI consensus sequences, while maintaining comparable accuracy (Fig. 4). With sequencing depth ten times deeper, both methods successfully identified all 43 mock 16S-23S rRNA operons in the three types of sequencing data (Supplementary Fig. 14a). Notably, the number of detected mock operons by kmerCon and miniCon was predominantly determined by sequencing accuracy. Both methods retrieved over 90% of operon copies in the PacBio CCS dataset, while less than half were identified in the two ONT datasets (Fig. 4a and Supplementary Fig. 14a). The UMI correction method demonstrated superior control of chimera, whereas corrected sequences based on sequence dissimilarity clusters contained 2-6% chimera, particularly in the PacBio CCS data.

**Fig. 4:**
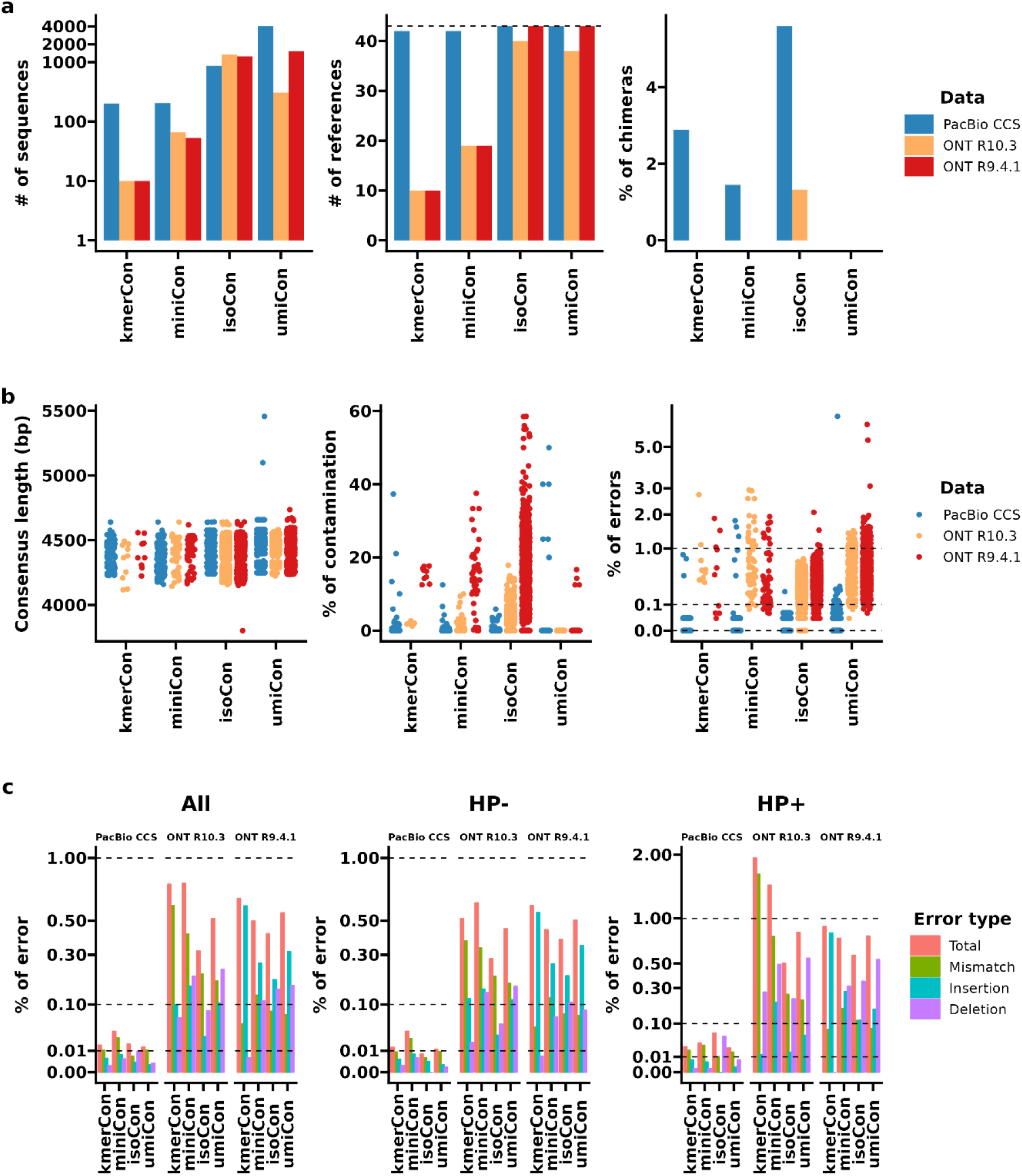
Statistics of various quality-controlled amplicon consensus sequences retrieved from the subsampled 100,000 reads tagged with unique molecular identifiers (UMIs). a, the number of consensus sequences and the identified reference sequences, and the chimera rate; b. the sequence length, the percentage of reads aligned to a reference sequence other than the primary class (contamination), and the error rate of each consensus sequence; c, the average error rate for mismatches, insertions, deletions, and total errors of retrieved consensus sequences, split by whether the error occurred inside (hp+) or outside (hp-) a homopolymer (hp) region. The corrected sequences are quality controlled with PCR artifacts removed and have recommended the minimum read coverage of 3×, 5×, 5× and 15 for the corrected sequences from umiCon, kmerCon, miniCon and isoCon, respectively. To enhance visualization, we applied a base-10 logarithmic scaling to the number of consensus sequences in sub-figure (a) and employed square root scaling for the error rates in sub-figures (b) and (c). The three datasets are colored by the sequencing platforms in sub-figure (a) and (b), and the bar colors in sub-figure (c) display different error categories. KmerCon, consensus calling on clustered sequences with Umapclust and Meshclust; miniCon, consensus calling on clustered sequences refined by overlap check; isoCon, consensus calling of detected isoforms by IsoCon on the clustered sequences; umiCon, consensuse calling on UMI bins.

All corrected sequences exhibited a consistent length distribution, approximately 4500 bp in size. UMI clusters displayed relatively low contamination, especially in the noisy ONT R9.4.1 data, while contamination rates were comparable across all consensus-calling approaches in the high-accuracy PacBio CCS data (Fig. 4b and Supplementary Fig. 14b). All error-correction strategies demonstrated remarkably low error rates below 0.1% in the accurate PacBio CCS reads after excluding a few rare clusters (Supplementary Fig. 15 and 16). Applying an acceptable cutoff of 5× for kmerCon and miniCon, and 15×for isoCon, these consensus correction approaches achieved overall error rates similar to those of 4×UMI corrections in the noisy ONT dataset (Fig. 4b). IsoCon consistently maintained error rates below 1%, which were 3-5 times higher than those of UMI corrections at the recommended coverage cutoff (Supplementary Fig. 14b). The error rates of isoCon and umiCon sequences decreased with read coverage, both converging to below 1% (Supplementary Fig. 15 and 16). With deeper sequencing, the error rate of umiCon fell below 0.1%, while the error rate of isoCon remained stable (Supplementary Fig. 16). At a shallow depth of 100,000 reads, the average overall error rate for 4×umiCon and 15×isoCon consensus sequences was 0.014% and 0.017% on PacBio CCS data, 0.55% and 0.42% on ONT R9.4.1 data, and 0.52% and 0.32% on ONT R10.3 data (see Fig. 4c and Supplementary Table 4), respectively. When the sequencing went ten times deeper, umiCon and isoCon sequences exhibited average error rates of 0.011% and 0.027% on PacBio CCS data, 0.071% and 0.401% on ONT R9.4.1 data, and 0.028% and 0.31% on ONT R10.3 data (Supplementary Fig. 14 and Supplementary Table 5), respectively.

The error types, including deletions, insertions, and mismatches, exhibited patterns specific to their data sources, with slight variations based on the consensus-calling approaches employed. The consensus sequences derived from PacBio CCS data displayed a distinct low error rate, showing no obvious differences in error types based on the presence of homopolymers. Mismatches predominantly contributed to errors, except for a notable presence of remaining deletion errors in homopolymer regions of the corrected isoCon sequences (Fig. 4c and Supplementary Fig. 14). Insertions in non-homopolymer regions were the most prevalent errors in ONT R9.4.1 consensus sequences, whereas mismatch errors were more common in ONT R10.3 corrected sequences (Supplementary Fig. 14). For ONT UMI consensus sequences, deletions in homopolymer regions and insertions in non-homopolymer regions were the major error sources (Fig. 4c). Similarly as in the PacBio CCS data, isoCon exhibited limited control over deletion errors in the ONT data (Supplementary Fig. 14 and Supplementary Table 5). The miniCon and kmerCon sequences however exhibited significantly higher levels of insertion errors in homopolymer regions when contrasted with other methods.

### ONT 16S rRNA gene amplicon sequencing of a synthetic mock microbiome

We followed a two-step PCR approach to prepare the ONT-based 16S rRNA gene amplicon library for microbiome profiling using metabarcoding^23^. We conducted a series of dilutions of ZymoBIOMICs mock DNA to test the multi-primer amplification library preparation and sequencing. At both sequencing depths of 10,000 and 100,000 reads per sample, all three consensus-calling methods effectively identified three primary OTU clusters, characterized by a length range from 900 to 1400 bp (Fig. 5a). These OTUs corresponded to the potential combinations of amplified hypervariable regions within the 16S rRNA operons, spanning from V1 and V3 to V8 and V9. Approximately half of the reads from each mock sample passed quality control for community profiling (Supplementary Fig. 17), and almost no OTUs were found in the negative controls (Fig. 5b). At the shallow sequencing depth of 10,000 reads per sample, the number of OTUs consistently remained below 46, which corresponds to the theoretical number of mock 16S rRNA operons. Nevertheless, with a tenfold increase in sequencing depth, kmerCon and miniCon generated 61 and 64 high-quality OTUs, respectively. Consistent with the results of the *in silico* test, isoCon split the 5-mer clusters and nearly tripled the theoretical number of OTUs (Supplementary Table 6). All eight mock strains were identified by all approaches at the species level (Fig. 5c). Some variations in relative abundance were observed, possibly due to PCR primer preferences, leading to an increase in *Limosilactobacillus fermentum* and a decrease in *Listeria monocytogenes* relative abundance in comparison to the theoretical composition. The retrieved OTUs demonstrated a high concordance with SILVA SSUs (Supplementary Figure 19). The dominant OTUs, i.e., the OTUs with minimal relative abundances of 0.25%, exhibited a species-level BLAST alignment identity of over 99% to the SILVA SSUs (Fig. 5d). These OTUs collectively accounted for over 97% of the total read counts (Supplementary Table 6). As sequencing depth grew, the profiling resolution of different mock DNA groups was improved with reduced inter-sample variance between technical replicates in the Bray Curtis distance metrices (Supplementary Fig. 18). The inter-group difference was eliminated when the phylogenetic relationship is integrated into beta-diversity analysis using the UniFrac dissimilarity metrics.

**Fig. 5:**
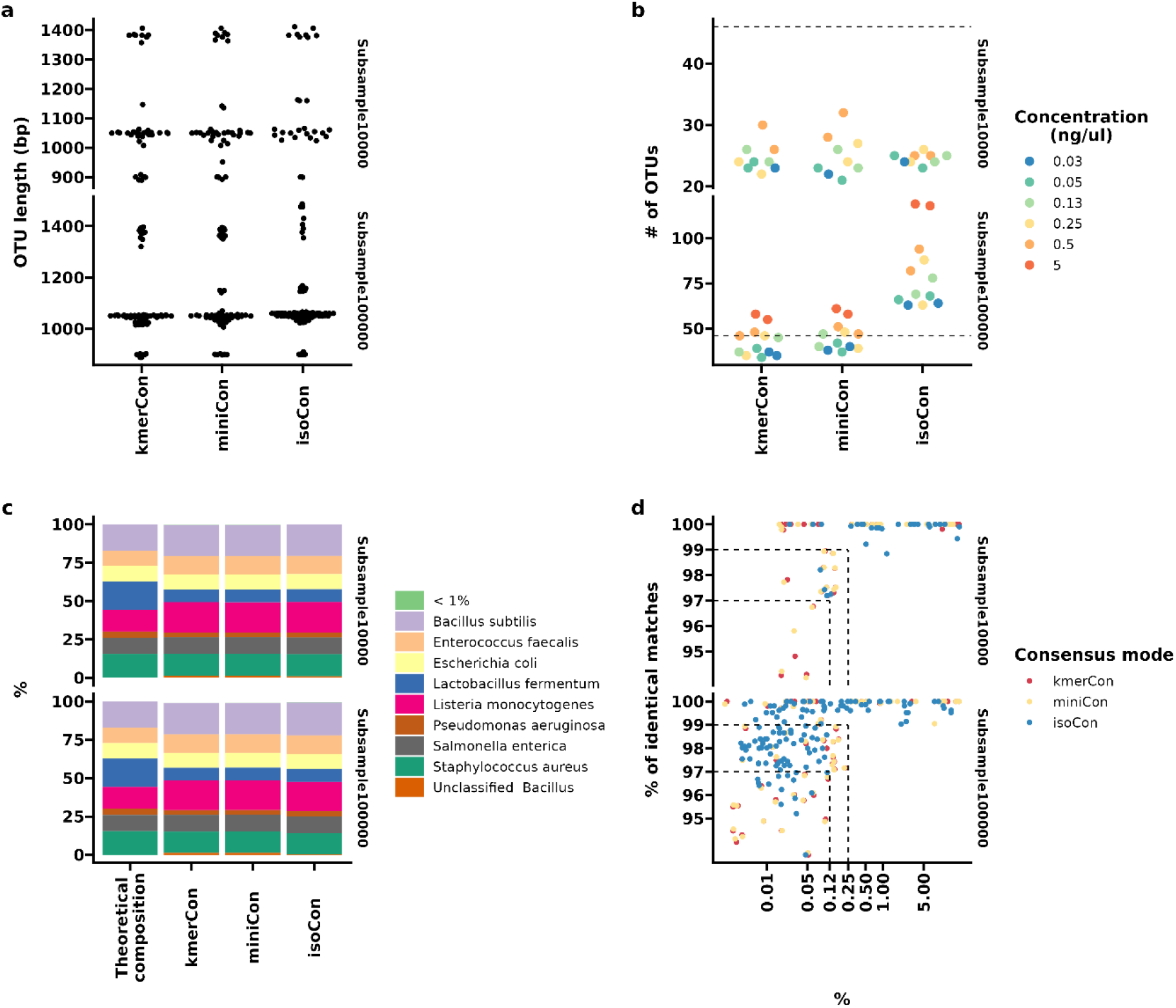
Near full-length 16S rRNA gene profiling of a mock dilution series through Oxford Nanopore sequencing. a, b, the length (a) and the number (b) of recovered OTUs by various OTU-picking approaches on mock serials with subsampled reads of 10,000 and 100,000 per sample; c, the species-level community profile and theoretical composition; d, the BLAST identity and relative abundance of picked OTUs. To enhance visualization, we applied a base-10 logarithmic scaling to the relative abundance of OTUs in sub-figure (d). The OTUs are colored by the mock concentration in the dilution series in the sub-figure (b) while the color in sub-figure (d) indicates the consensus calling modes. Taxa with a mean abundance of less than 1% are marked in sub-figure (c) as “< 1%”. KmerCon, community profile based on clustered sequences with UMAPclust and Meshclust; miniCon, community profile based on clustered sequences refined with overlap check; isoCon, community profile based on detected isoforms by IsoCon on the clustered sequences.

### Real-world application in human vaginal microbiomes

To assess LACA’s performance in a real-world application, we utilized a public ONT 16S rRNA gene amplicon dataset^10^ of 12 human vaginal microbiomes composed of six controls and six patients with diagnosed bacterial vaginosis, and employed the read-classification approach Emu^10^ as a comparison. These approaches consistently identified the dominant bacteria on the vaginal swaps at genus (Supplementary Fig. 20) and species level, except that some species-level signals were conservatively annotated to the genus level by LACA given the use of local common ancestor (LCA) algorithm (Supplementary Fig. 21). In comparison to a normal vaginal microbiome, all methods revealed a significant increase of non-lactobacilli anaerobes, such as *Aerococcus*, *Prevotella*, and *Megasphaera*, in samples from patients with bacterial vaginosis (Supplementary Fig. 21 and 22). The difference between these two sample types was more pronounced in the weighted UniFrac metrics, owing to the phylogenetic information preserved in OTUs (Fig. 6). Emu identified approximately 3 times more taxonomic features at the species level and 0.4 times more at the genus level compared to LACA (Supplementary Fig. 23). KmerCon identified the most taxonomic traits at the genus and species levels among the LACA profiles, followed by miniCon and isoCon. The shared features in the LACA and Emu profiles showed a strong positive correlation in relative abundance, with an average Pearson’s correlation coefficient above 0.95 (*p* < 0.05 Supplementary Fig. 24). Although isoCon identified the fewest taxonomic features, the genus-and species-level abundances were most correlated with the estimated abundance through re-mapping reads against the OTU sequences (Supplementary Fig. 25). The correlation between clustering-based and alignment-based abundance declined at the OTU level while the Spearman rank test still indicated a strong correlation. Since LACA OTUs were annotated by LCA algorithm based on BLAST hits, we searched the hit table for the uncovered taxonomic features by LACA. Among all taxonomic features identified by Emu, over 70% and 65% of species and genus annotations were found in the BLAST hit table against SILVA SSUs with a minimum alignment identity above 97% and 99%, respectively (Supplementary Table 7). The relative abundance of the uncovered genus and species was below 0.1%, except for *Criibacterium bergeronii* and the *Rikenellaceae* RC9 gut group (Supplementary Table 8). In BLAST search against the curated EzBioCloud database for 16S rRNA gene sequences, we found that the *Peptostreptococcaceae* OTUs showed over 99.1% identity to the 16S rRNA gene of the *Criibacterium bergeronii* strain CCRI-22567 in the best hit (Supplementary Table 9). LACA identified the *Rikenellaceae* RC9 gut group, but the relevant OTUs were excluded in analysis due to a low alignment identity of around 90% among all hits against SILVA SSUs. The LCA assignment of one OTU sequence was influenced by the redundant and un-curated species-level taxonomy in the SILVA database and the homogeneity of 16S rRNA gene copies among closely related species. For example, the lactobacilli OTUs were not annotated as *Lactobacillus iners* in the control vaginal samples (Supplementary Fig. 21) although BLAST hits suggested over 99% identity (Supplementary Table 8). Meanwhile, Emu missed some vaginal members, including nine species and three genus features, such as *Variovorax* and *Solobacterium*, relative to the LACA approach (Supplementary Table 10). The relative abundances of the uncovered features were below 0.1%, except for *Lactobacillus rhamnosus* and *Lactobacillus fermentum*, which accounted for 0.2% and 0.7%, respectively.

**Figure 6.**
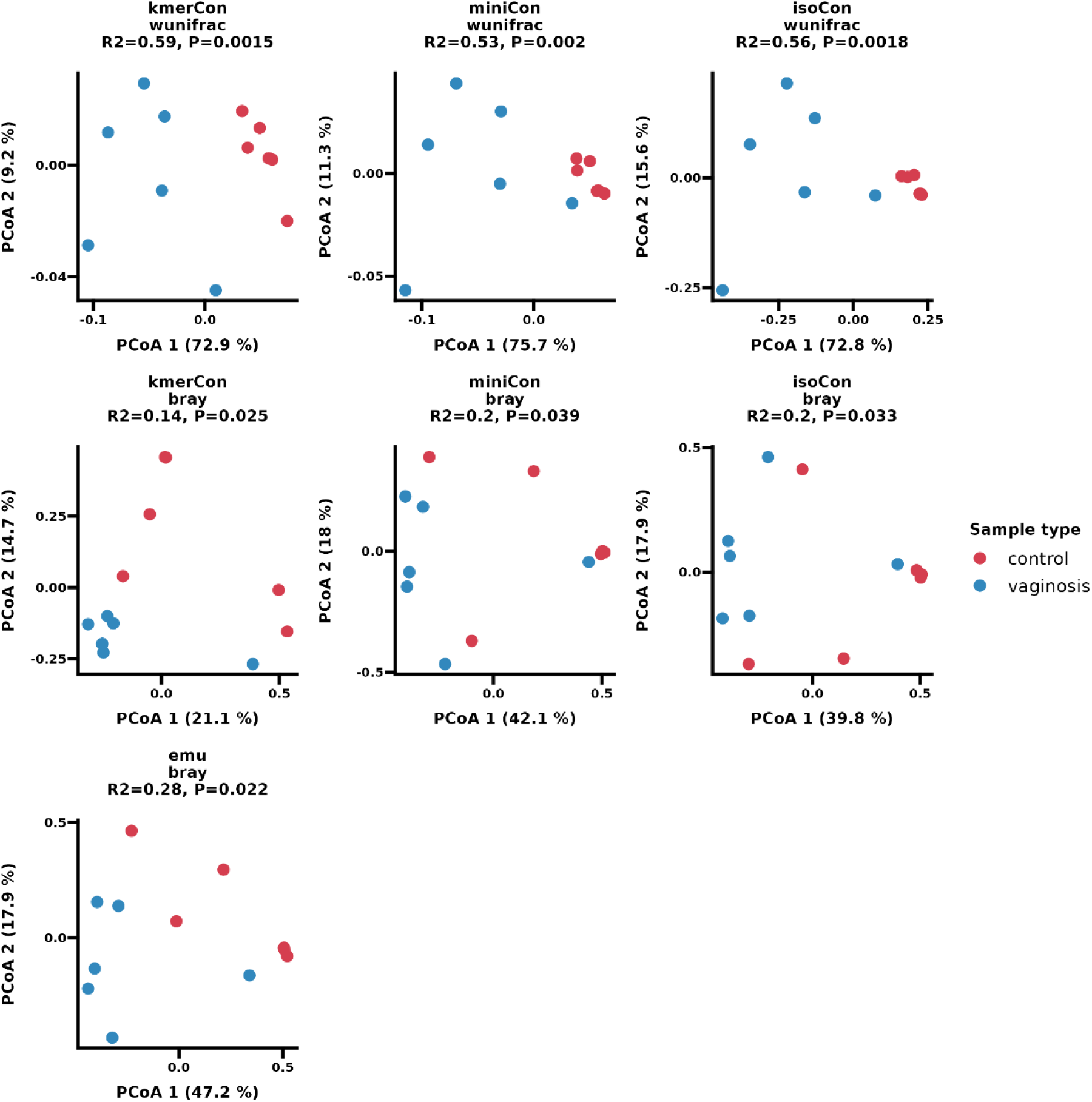
Principal coordinate analysis (PCoA) plots of weighted UniFrac and Bray Curtis dissimilarity using various profiling approaches on the human vaginal samples. The points are colored by the sample phenotype. Emu, read classification profile with Emu; kmerCon, community profile based on clustered sequences with UMAPclust and Meshclust; miniCon, community profile based on clustered sequences refined with overlap check; isoCon, community profile based on detected isoforms by IsoCon on the clustered sequences.

## Discussion

Utilizing a *de novo* OTU picking strategy, LACA offers a full microbiome analysis suite tailored for long-read amplicon analysis, spanning from demultiplexing to taxonomic quantification. Unlike read classification approaches, *de novo* OTU picking enables novel species detection without reference bias, and it encodes phylogenetic information in nucleotides, offering feasibility in microbiome meta-analysis.

The variable sequencing error rate poses a challenge for OTU picking from long-read amplicons. While these sequencing errors are generally randomly distributed^24,25^, they can obscure true variants in highly similar regions. To address this, LACA benchmarked multiple alignment-free and align-based methods, including the use of short k-mers^13,14^ or error-aware clustering with long minizers^11^, or iterative clustering through alignment^15^. In our test, we observed the clustering choice was determined by the sequence divergence of the target region and the basecalling accuracy of the applied technologies. To avoid the unnecessary computational cost of pairwise alignment, LACA first clusters sequences with alignment-free approaches. Among these, UMAPclust, incorporating 5-mer frequency information, displayed superior efficiency in clustering long 16S rRNA gene amplicons with few false positives (Fig.1). In combination with Meshclust^19^, UMAPclust accurately identified ONT R9.4.1 amplicons of 16S rRNA genes with minimal sequence divergence of 5%. Its resolution improved to 2% and 1% for ONT R10.4.1 and high-accuracy ONT Duplex and PacBio CCS data, respectively (Fig. 2). These varying results highlight the trade-off between errors and variants inherent in the methodology. UMAPclust relies on the use of short 5-mers, which are highly sensitive to point variations, making it effective in many cases. However, it is vulnerable to producing distorted distributions when insertions and deletions are introduced^26^. It is a significant concern in long-read technologies since indels serve as common sources of errors ^27^. Thereby its performance significantly improved in the latest ONT R10.4.1 flowcells using dual-reader nanopore to reduce indel calling errors in the homopolymer region. The alignment-free clusters could be further refined through alignment-based approaches. Combining isONcorrect and isoCon improved cluster purity and extended identifiable sequence divergence to 1% for ONT R9.4.1 data and below 1% for ONT R10.4.1, high-accuracy Duplex and PacBio CCS data. With this refinement, the error rate of consensus sequences was controlled below 1% for ONT R9.4.1 data, 0.2% for ONT R10.4.1 and 0.1% for high-accuracy long-read data (Supplementary Fig. 11). Given that acceptable species-level resolution for 16S rRNA genes falls within the range of 97-99%, these corrected sequences have effectively preserved phylogenetic information for precise microbiome analysis. In a multi-primer amplicon library designed for comprehensive sequencing of the 16S rRNA gene, we confirmed that the selected OTUs accurately delineated eight mixed mock strains at the species level using noisy ONT R9.4.1 data (Fig. 5).

To mitigate long-read sequencing errors, specialized library preparation methods such as UMI^5^ and rolling circle amplification^28^ have been implemented to ensure molecular-level precision. Nonetheless, molecular-level correction requires sufficient sequencing depth for each individual molecule. In the case of sequencing multi-copy gene families like rRNA genes, the use of UMI tags can result in redundant tagging of identical copies originating from a single template. This leads to a significant increase in sequencing costs, typically requiring optimal dilution strategies to strike a balance between adequate sampling and efficient utilization of sequencing throughput. Determining the appropriate dilution level is challenging, particularly when dealing with unfamiliar ecosystems. In shallow sequencing, corrections through sequence similarity clustering demonstrated comparable or even superior sensitivity and error control compared to 4×UMI corrections to identify highly similar 16-23S rRNA gene operons (Fig. 3a). Despite no further error reduction with increased sequencing depth, most corrected sequences maintained acceptable error rates below 1% for species-level alignment. On high-fidelity PacBio CCS data, the UMI correction and clustering-based approaches exhibited a similar ability to reduce errors to below 0.1% with adequate coverage cutoffs. The ONT Duplex kit with the R10.4.1 flowcell is expected to offer single-molecule resolution with an average error rate of Q30. This shall further broaden the utility of clustering-based corrections for long amplicon analysis.

The OTUs generated through *de novo* clustering are anticipated to faithfully retain the phylogenetic information originating from the profiled samples. Based on UniFrac distance metrics, we found that these long OTUs effectively contribute to characterizing sample phenotypes, even when working with the prevailing ONT 9.4.1 flowcell whose error rate can range from 5-15%. In a beta-diversity analysis of a synthetic mock community, significant differences in Bray Curtis dissimilarity were observed due to variations in PCR template concentrations. These differences became more pronounced with deeper sequencing, while disparities arising from technical replicates diminished. Stochastic fluctuations in priming efficiency, a consequence of varying template concentrations, likely contributed to this phenomenon^29,30^. However, the incorporation of evolutionary information through UniFrac distance metrics effectively mitigated these differences. Another noteworthy example is the characterization of vaginal microbiomes. Healthy vaginal microbiomes are typically dominated by lactobacilli^31^. The phylogenetic resemblance in the amplification region led to improved clustering of these samples when evaluated using the weighted UniFrac distance metric. This phylogenetic weighted clustering facilitated a clear separation between control samples and those associated with vaginosis, surpassing even the Emu profile, which incorporated more taxonomic features based on read classification.

While clustering-based approaches exhibited decreased sensitivity in detecting rare species compared to read classification tools such as Emu, LACA effectively identified all dominant microorganisms (> 0.1%) at both the genus and species levels, demonstrating strong agreement between the two methods. *De novo* picked OTUs require a minimum cluster size to distinguish the true biological unit from spurious sequences, rendering them predisposed to potentially overlooking subtle signals from rare species. On the other hand, augmented sensitivity of profiling with read classification carries a higher risk of generating false positives. The original study^10^ notes escalated false-positive counts of Emu in characterizing a synthetic community relative to NanoCLUST^14^, despite Emu using an expectation-maximization algorithm to iteratively refine species-level abundance. This discrepancy could be attributed to mapping errors introduced during the initial alignment of long, noisy reads against highly similar but redundant gene sets, such as rRNA operons. In our analysis, we encountered situations where the LCA taxonomy for certain OTUs could not be definitively resolved at the species level, even though they exhibited over 99% BLAST identity to reference sequences (as observed with *Lactobacillus iners* and *Criibacterium bergeronii* in vaginal microbiomes). The underlying causes of this issue are related to the sequence homogeneity of closely related species in the amplification region and a substantial portion (∼72%) of the undetermined species labels in the SILVA database^32^. This underscores the biological limitations of the 16S rRNA gene for achieving a comprehensive taxonomic profile at the species level. Analogous findings have been reported in efforts to utilize the complete 16S-23S rRNA gene operon for species-level assignments^5^, although improved species-level resolution have been observed with the increase in amplicon length.

LACA provides a versatile, reproducible, and scalable workflow for *de novo* OTU analysis of lengthy noisy amplicon data. In our assessment, the clustering-based OTUs effectively preserved the essential phylogenetic information for long amplicon microbiome analysis, with an acceptable trade-off in accuracy. Although we benchmarked LACA with rRNA operons for microbiome analysis, these clustering-based corrections can also be applied to other types of amplicon data within a discernible range. Furthermore, LACA’s flexibility extends to its configuration file, making it easy to adapt to novel models and flowcells as sequencing technologies continue to evolve.

## Methods

### Data generation

#### Long amplicon simulation on SILVA SSU rRNA sequences

The whole simulation process was accomplished with the simulation module in LACA workflow. In short, the reference sequences were selected according to the sequence divergence requirements (of 5-10%, 3-5%, 2-3%, 1-2%, and 0-1%) through two rounds of MMseqs2^33^ clustering of SILVA^34^ 138.1 SSU rRNA sequences and were used as reference sequences for read simulation by Badread^35^. For each selected reference set, ten sequences were picked and had the same length of 1522 bp after being clustered by minimal and maximal sequence identity. The relative abundance of each reference sequence was set to equal (even10) or skewed with two abundant sequences 10 times as much as the rest (skew10). The sequence identity of ONT R9.4.1 and ONT R10.4.1 reads followed a beta distribution with mean, max, and standard deviation of 87.5%, 97.5% and 5% using the error and quality model of “nanopore2020”, and 95%, 99%, and 2.5% for the “nanopore2023” model, respectively. The simulation identity of high-accuracy ONT duplex and PacBio HiFi reads followed a normal quality score distribution with a mean and standard deviation of 20, 4 using the error models “nanopore2023” and “pacbio2016,” respectively. This led to four sets of long-read sequencing data, namely “ONT2020”, “ONT2023”, “ONT Duplex” and “PacBio CCS” in the text. By default, the simulated reads included 1% of chimeras and were appended with the ligation adapters of 5’-AATGTACTTCGTTCAGTTACGTATTGCT-3’ and 5’-GCAATACGTAACTGAACGAAGT-3’ on both ends of either strand. The i*n silico* read coverage was set to 40×, 60×, 80×, 100×, 200×, 400×, respectively. Thus, the simulation process resulted in 120 independent read files (from various sequencing platforms and depth), for the following performance test.

#### Near full-length 16S rRNA amplicon sequencing on diluted mock serials DNA source

The ZymoBIOMICS microbial community DNA standard (D6306, Zymo Research) was used as mock DNA for library preparation. This commercial mock contains DNA from ten different microorganisms, including eight bacteria and two yeasts: *Pseudomonas aeruginosa*, *Escherichia coli*, *Salmonella enterica*, *Limosilactobacillus fermentum*, *Enterococcus faecalis*, *Staphylococcus aureus*, *Listeria monocytogenes*, *Bacillus subtilis*, *Saccharomyces cerevisiae*, and *Cryptococcus neoformans*. The original mock DNA was diluted in series with sterile water, resulting in samples with the mock DNA concentrations of 5, 0.5, 0.25, 0.13, 0.05, and 0.03 ng/µL, respectively. The DNA concentration was measured with Qubit 1× dsDNA HS assay kit (Invitrogen, Thermo Fisher Scientific).

#### Target gene amplification and metabarcoding

The library preparation followed a published protocol^23^ for ONT sequencing of near full-length 16S rRNA amplicons with minor modification. In short, near full-length 16S rRNA gene was amplified with a protocol involving multiple forward and reverse primers: 27Fa/b (5’- GTCTCGTGGG CTCGGNNNNN NNNNNNNNNN AGAGTTTGAT YMTGGCTYAG -3’, 5’- GTCTCGTGGG CTCGGNNNNN NNNNNNNNNN AGGGTTCGAT TCTGGCTCAG -3’), 338Fa/b (5’- GTCTCGTGGG CTCGGNNNNN NNNNNNNNNN ACWCCTACGG GWGGCAGCAG -3’, 5’- GTCTCGTGGG CTCGGNNNNN NNNNNNNNNN GACTCCTAC GGGAGGCWG CAG -3’), 1391R (5’- GTCTCGTGGG CTCGGNNNNN NNNNNNNNNN GACGGGCGGT GTGTRCA -3’) and 1540R (5’- GTCTCGTGGG CTCGGNNNNN NNNNNNNNNN TACGGYTACC TTGTTACGACT-3’). First PCR conditions were as follows: 95°C for 5 min, 2 cycles of 95°C for 20 s, 48°C for 30 s, 65°C for 10 s, 72°C for 45 s, and a final extension at 72°C for 4 min. A second PCR step was done to barcode the first PCR products with the following conditions: 95°C for 2 min followed by 33 cycles of 95°C for 20 s, 55°C for 20 s, 72°C for 40 s, and a final extension at 72°C for 4 min. PCR products were cleaned up using AMPure XP beads (Beckman Coulter Genomic, CA, USA) after each PCR step. After validation with agarose gel electrophoresis, the final PCR products were pooled for ONT sequencing.

#### ONT sequencing with barcoded amplicons

The ONT sequencing library was constructed according to the ligation sequencing kit SQK-LSK109 protocol. The library was loaded on R9.4.1 flow cell and sequenced with GridIONX5 platform (Oxford Nanopore Technologies, Oxford, UK). Sequencing data was collected by MinKnow (v21.05) for Guppy (v5.0.11) basecalling in high-accuracy mode.

### Long amplicon analysis

#### Clustering quality comparison of alignment-free approaches on simulated SILVA amplicons

To assess clustering performance of alignment-free approaches, LACA was adapted to treat the 20 simulated SILVA amplicon read files (under a fixed and adequate sequencing depth of 200×) as independent samples and consensus calling was performed without sample pooling to avoid cross-contamination from the other simulation scenarios e.g., the different sequencing platform. The ligation adapters were treated as primers by LACA, and the *in silico* reads were checked with Cutadapt^36^ for the linked primer pattern of 5’- AATGTACTTCGTTCAGTTACGTATTGCT… GCAATACGTAACTGAACGAAGT -3’) with a maximal error rate of 0.2 and minimal overlaps of 6 bases. The primer sequences were retained in the checked reads and extra bases were trimmed. Chimeric reads were filtered by yacrd^37^ with minimal read coverage of 0.4 and the minimal coverage was set to 4 and 3 for ONT and PacBio CCS amplicons, respectively. The read length range was set to 800-2000 bp and low-quality reads were discarded if the average quality score was below 7. The quality-controlled reads were sent for alignment-free clustering with isONclust^11^, UMAPclust and Meshclust^19^.

We benchmarked the effect of various combinations of these alignment-free approaches and the Meshclust^19^ thresholds on the clustering performance of the SILVA amplicons from the four types of long-read sequencing data. To determine appropriate Meshclust^19^ thresholds for respective long read data, we used LACA in the “clust” mode only with Meshclust^19^ using a set of the identity scores of 0.7, 0.75, 0.8, 0.85 and “auto” for the ONT2020 reads, 0.8, 0.85, 0.9, 0.95 and “auto” for the ONT2023 reads and 0.85, 0.9, 0.95 and “auto” for the ONT Duplex and PacBio CCS reads. In the clustering benchmark of various approaches and the combinations, the estimated identity score by Meshclust^19^ was applied for all four data types. And LACA was performed in “clust” mode, employing isONclust^11^, UMAPclust, or a sequential combination of both, culminating in the final step of Meshclust^19^ clustering. For isONclust^11^, ONT reads were clustered with the recommend parameters of “-k 13 -w 20” and PacBio CCS reads were clustered with the flag “-k 15 -w 50”. The UMAPclust clustering stuck to the settings of NanoCLUST^14^ with HDBSCAN^38^ clustering on the two-dimensional UMAP^39^ graph of the canonical 5-mer frequency of these amplicon reads. The UMAP^39^ adopted similar parameters with “n_neighbors=15, min_dist=0.1, metric=cosine” and the HDSCAN^38^ parameters were set as “min_bin_size=10, min_samples=10, epsilon=0.5”.

#### Consensus calling evaluation on the simulated SILVA amplicons from various sequencing depth

To assess performance of kmerCon, miniCon, isoCon, LACA was adapted to treat the 120 simulated SILVA amplicon read files as independent samples for consensus calling to avoid cross-contamination from the other simulation scenarios. The quality control process stuck to the same procedures as that in the above section “Clustering quality comparison of alignment-free approaches on simulated SILVA amplicons”. The passed reads were first clustered by the alignment-free approaches with UMAPclust and Meshclust^19^, and the three types of consensus calling were conducted within the clustered reads. Given increasing *O(n^2^)* of pairwise alignments with cluster size, miniCon was conducted in batches with a maximum size of 5,000 reads per batch. And the minimal fraction of maximum score for a miniCon cluster was set to 0.75 and 0.85 for ONT2020 and the rest read types. In the isoCon mode, read correction by isONcorrect^16^ was not enabled for high-accuracy ONT Duplex and PacBio CCS data to maximally preserve the variant signals. Each consensus sequence had at least 20 supporting reads and had two rounds of Racon^22^ polishing. Extra one round of Medaka polishing was introduced to produce the final ONT consensus sequences. The “r941_min_hac_g507” consensus model was used for the ONT2020 amplicons while the “r1041_e82_400bps_hac_v4.2.0” consensus model was adopted for the ONT2023 and ONT Duplex amplicons. For ONT Duplex and PacBio CCS data, we also conducted consensus calling within the clusters generated by isONclust^11^ and UMAPclust. This was done under the loose supervision by Meshclust^19^ using an identity score of 0.5, which maximally preserved the clusters in the initial two round of clustering.

#### Consensus calling evaluation on UMI-tagged long amplicons

We utilized the PacBio CCS, ONT R9.4.1 and R10.3 UMI-tagged amplicon datasets^5^ of the ZymoBIOMICS mock and subsampled each to 100,000 and 1,000,000 reads. The consensus sequences were extracted with LACA according to the sequencing platform and clustering approaches. For kmerCon, miniCon and isoCon, we followed similar quality control and consensus calling procedures in the section “Clustering quality comparison of alignment-free approaches on simulated SILVA amplicons”. The forward (5’- AGRGTTYGATYMTGGCTCAG-3’) and reverse primer (5’- CGACATCGAGGTGCCAAAC-3’) were checked with outside bases trimmed, and chimeric reads were excluded. To keep consistent with original publication^5^, the allowed read length range was set from 3500 to 6000 bp and the minimal average quality score was 7. Then UMAPclust and Meshclust^19^ were used for alignment-free clustering ONT reads, and the “min_bin_size=50, min_samples=50” was used for HDBSCAN in UMAPclust and the estimated identity score was used for Meshclust^19^. The PacBio CCS reads were clustered by isONclust^11^, UMAPclust and Meshclust^19^ in a sequential order and the Meshclust^19^ identity score was 0.5. Given increasing *O(n^2^)* of pairwise alignments with cluster size, miniCon was conducted in batches with a maximum size of 5,000 reads per batch. And the minimal fraction of the maximum score for a miniCon cluster was set to 0.85. For isoCon, read correction by isONcorrect^16^ was enabled for ONT data, and consensus calling was performed in batches (with a maximum size of 20,000 reads per batch). Each clustering-based consensus sequence had at least 3 supporting reads. The polishing phase by Racon^22^ or Medaka stuck to the same settings as that in UMI-based corrections^5^. For umiCon, the consensus sequences were extracted using the 36-bp UMIs (5’-NNNYRNNNYR NNNYRNNNNN NYRNNNYRNN NYRNNN-3’) following the published procedures^5^ with open-source software after a simple read length check. The existence of primers was checked and retained on the corrected sequences and extra bases were trimmed.

#### Full-length 16S rRNA gene amplicon analysis of a diluted mock series

Demultiplexing was accomplished using Guppy (implemented in LACA) with custom barcodes. To evaluate the LACA performance on the mock samples, the demultiplexed reads were subsampled to 10,000 (subsample 10,000) and 100,000 (subsample 100,000) per sample if possible. Similarly, we followed the same consensus calling procedures in the section “Consensus calling evaluation on UMI-tagged long amplicons” unless otherwise specified. The spurious and chimeric amplicon reads were filtered if the sequence length exceeded the range of 800 to 1600 bp. Given the multiple-primer strategies in amplification, reads were retained if they contained any possible combinations of linked patterns of the forward and reverse primers. The remaining reads were first pooled together for alignment-free clustering of UMAPclust and Meshclust^19^ (with a minimal identity score of 0.9), followed by consensus calling with kmerCon, miniCon and isoCon. Each consensus sequence had at least 10 supporting reads and two rounds of Racon^22^ polishing followed by one round of Medaka polishing using the “r941_min_hac_g507” consensus model. The OTUs were the representative sequences picked through MMseqs2^33^ clustering with an alignment coverage above 0.9 and a sequence identity above 0.99. The OTU matrix was constructed according to the preserved sequence ID information in the OTU clustering and demultiplexing. The phylogenetic tree was built with FastTree^40^ implemented in QIIME 2^17^ and the OTU taxonomy was determined by QIIME 2^17^ classify-consensus-blast against the SILVA^34^ 138.1 SSUs. By default, the LCA taxonomy was taken for each OTU sequence based on the agreement among ten BLAST hits.

#### Full-length 16S rRNA gene amplicon analysis of human vaginal microbiomes

To evaluate LACA performance in real-world applications, we re-utilized the public human vaginal dataset in the Emu^10^ paper. The LACA settings similar as that in the previous section “Full-length 16S rRNA gene amplicon analysis of a diluted mock series” unless otherwise specified. According to the chosen primer sets (27F: AGAGTTTGATCMTGGCTCAG and 1492R: CGGTTACCTTGTTACGACTT), the expected amplification range was 1300 to 1700 bp. We used UMAPclust and Meshclust^19^ (with the estimated identity scores) for initial alignment-free clustering, followed by consensus calling with kmerCon, miniCon and isoCon. And each OTU sets were picked from the respective corrected sequence sets with an alignment coverage above 0.99 and a sequence identity above 0.99. To maintain consistency with the Emu profile, the respective OTU sets were picked without sample pooling. To compare with the OTU matrix based on clustering information, we mapped quality-controlled reads back to the OTU sequences, generating an alignment-based count matrix. The Emu profiles of each sample were processed with our open-source workflow NART (https://github.com/yanhui09/nart) and merged into one matrix by the detected taxonomic features. Similarly, the primer sequences were retained for Emu alignment and the sequencing data were quality controlled as that in the LACA process. The SILVA 138.1 database was adopted to determine the taxonomic assignments in all profiling approaches.

### Data analysis

#### Clustering evaluation on *in silico* data

The clustering quality was assessed by examining the number of clusters and two external metrics: purity and NMI. The external metrics were calculated using the true class labels of the simulated data as a reference. The clustering purity measures how homogeneous a cluster is with respect to a single class^41^, and is defined in Eq. (1) as below:

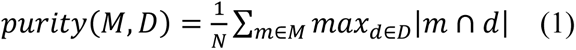

where *M* is the set of clusters *m*, *D* is the set of classes *d* and *N* is the number of clustered reads. NMI measures the similarity between the clustering results and the ground truth and trades off the clustering quality against the number of clusters^41^, and is defined in Eq. (2) as below:

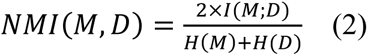

where *M* is the set of the clusters, *D* is the set of classes, *I* is the mutual information as defined in Eq. (3), and *H* is entropy as defined in Eq. (4).

Given *p(m)*, *p(d)* and *p(m∩d)* are the probabilities of a read being in cluster *m* of the set *M*, class *d* of the set *D*, in the intersection of *m* and *d*, *I(M;D)* is defined as below:

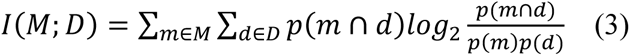

Given *p_i_* is the probability of label *i* in the set *M*, *H(M)* is defined as below:

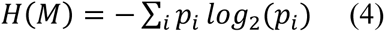

The clustering purity was calculated by custom R code and NMI by R package aricode^42^ as stated in the “Code Availability” section.

#### Alignment validation with simulation reference sequences

The consensus sequences were split by the sequencing depth, simulation reference set, and consensus-calling methods after the adapter sequences were trimmed from both sides with Cutadapt^36^. The split sequences were aligned against respective reference sequences at the base level with Minimap2^43^ under the preset of asm5, asm10, and asm20, and only primary alignments were kept. Given the sequence redundancy of SILVA SSUs, the frequent minimizers occurring more than 10,000 times were ignored in Minimap2^43^ mapping. The alignment identity was calculated by dividing the total bases, including gaps in the mapping, by the number of matching bases. The query (consensus) sequence cover was calculated by dividing the number of matches by the length of the query sequence. The target (reference) sequence cover was calculated by dividing the number of matches by the length of the target sequence.

#### Error profiling of consensus sequences from UMI-tagged data

The derived consensus sequences were validated using the script “longread qc_pipeline” with the curated 16S-23S rRNA operon ZymoBIOMICS reference and SILVA^34^ 132 SSURef Nr99 reference as in publication^5^. The provided R script “validation functions.R” was used to compile the generated results for error profiling and quality evaluation. Briefly, the error types (mismatch, deletion, insertion) and the relative positions of the errors were determined with respect to the reference sequence. These errors were then categorized as being within homopolymer regions (hp+) or not (hp-). Accordingly^5^, the error information combined with other data (such as the number and length of consensus sequences, cluster contamination, ZymoBIOMICS reference-based taxonomy, SILVA taxonomy, and chimera detection) was calculated before and after the PCR artifacts were removed. The consensus sequences were flagged as PCR artifacts if the errors are evenly distributed and if the operon SSU part has a better match with SILVA SSU than the ZymoBIOMICS reference. The remaining PacBio homopolymer artefacts were manually detected and flagged in the corrected sequences. The function “lu_artifact_plot” in the provided R script was used to identify artifacts, using cluster size intervals of 3 (when possible) from 1 to 60 and larger than 60 for UMI clusters. The intervals for other types of clustering methods were set using quantiles of 0%, 25%, 50%, and 75% and greater than 75% due to the significant decrease in the number of clusters. The cluster contamination was estimated by the percentage of the clustered reads assigned to different ZymoBIOMICS reference-based taxa except for the most common taxonomic class. The chimeras were identified if called by UCHIME2^44^ with the curated ZymoBIOMICS reference and if errors tended to occur more frequently on either end of a sequence.

#### Community analysis of the diluted mock serials and human vaginal microbiomes

The clustering-based count matrices were used for community analysis unless otherwise clarified. We integrated the count matrix, phylogenetic tree file, and LCA taxonomy file for community analysis with R package phyloseq^45^. The result generated by each approach was analyzed independently, and high-quality OTUs were retained for community analysis with at least one BLAST hit showing alignment identity above 97% to the SILVA SSUs. Hellinger transformation was applied on the raw count matrix for dissimilarity metrics computation in the vaginal dataset while the raw count matrix was used for the subsampled synthetic mock dataset. Bray Curtis and weighted UniFrac dissimilarity metrics were calculated in comparison for PERMANOVA test using R package vegan^46^. The BLAST hits for LCA taxonomy were collected to evaluate the alignment identity and cover. The analysis script is provided in the “Code Availability” section.

#### Intersection visualization and correlation analysis of taxonomic features

The taxonomic features from various profiles were extracted at species and genus level and the intersections between matrix were visualized in UpSet^47^ plot. Spearman’s rank and Pearson correlation analysis of the relative abundance of shared taxonomic features were conducted between various LACA profiles and Emu output, and the rare features present among less than 30% of samples were excluded. The same procedures were applied for the correlation analysis between the count matrices constructed by the clustering-based and mapping-based approach. The analysis script is provided in the “Code Availability” section.

## Supporting information

Supplementary Figures and Table titles

## Data availability

The near full-length amplicon Nanopore sequencing results of the ZymoBIOMICS microbial community standard D6300 were deposited in the NCBI Sequence Read Archive as Bioproject PRJNA1036680. Public data used in this investigation include the ZymoBIOMICS microbial community standard D6300 UMI-tagged sequencing data from the European Nucleotide Archive under the project PRJEB32674, the 12 vaginal samples in real-world application from Sequence Read Achieve under the project PRJNA723982 and the SILVA 138.1 SSURef Nr99 database (https://www.arb-silva.de/).

## Code availability

LACA is open-source and available at https://github.com/yanhui09/laca. The LACA v0.3.0 was used to process the long amplicon reads in this paper. NART is open-source and available at https://github.com/yanhui09/nart. The data analysis scripts can be accessed at https://github.com/yanhui09/laca_archive.git.

## Acknowledgements

We acknowledge the high-performance computing resources provided by Danish national supercomputer for life sciences (Computerome, https://computerome.dk).

