## Supplementary Figures and Table titles for "*De novo* clustering of long-read amplicons improves phylogenetic insight into microbiome data"

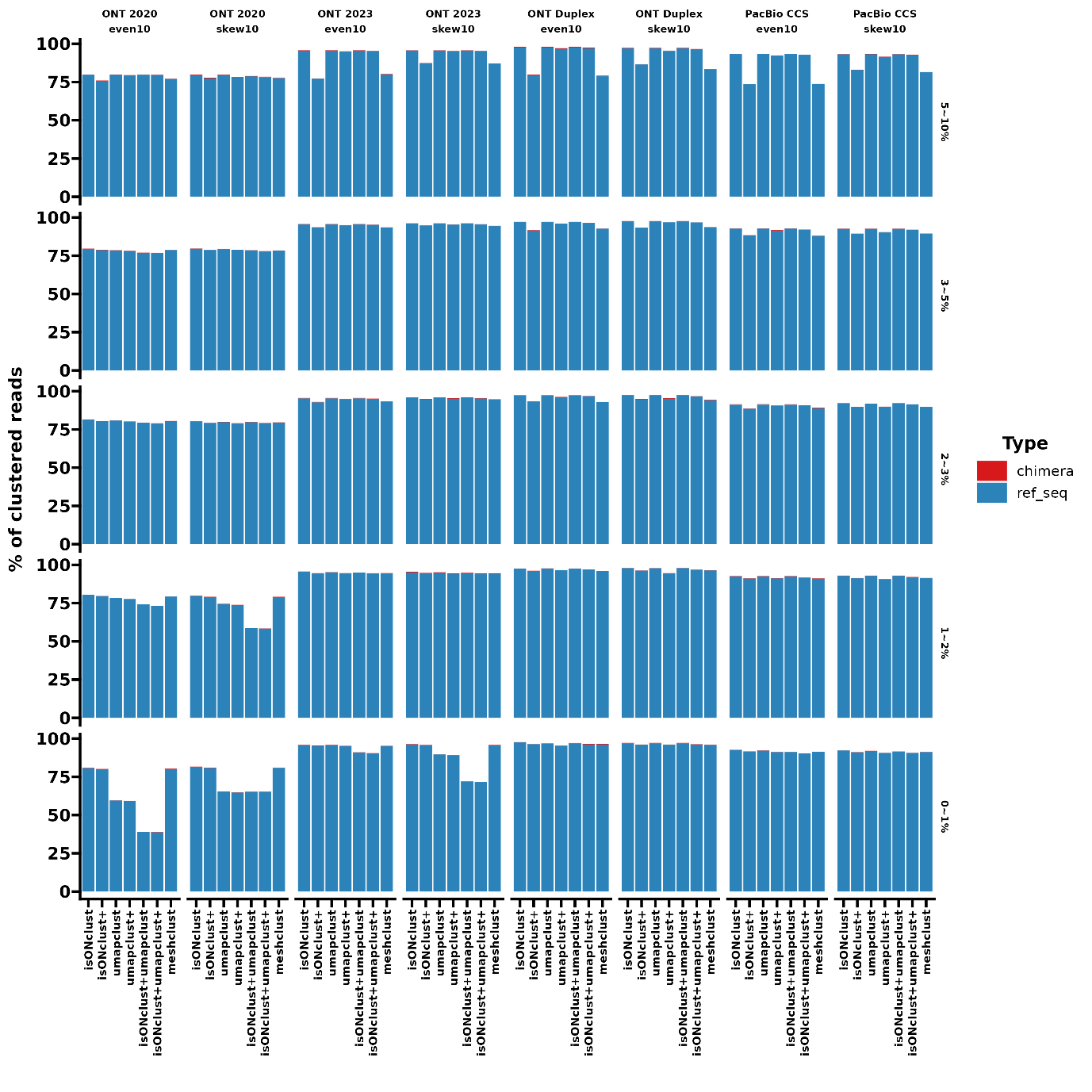

Supplementary Fig. 1. The portion of clustered reads with various alignment-free approaches on the SILVA amplicons under the *in silico* sequencing depth of 200×. The portion is colored by the simulated read types. The reference sequences are chosen from SILVA small subunit sequences with the sequence divergence of 5-10%, 3-5%, 2-3% 1-2%, and 0-1%. KmerCon, consensus calling on clustered sequences with UMAPclust and Meshclust; miniCon, consensus calling on clustered sequences refined by overlap check; isoCon, consensus calling of detected isoforms by IsoCon on the clustered sequences. Even10 refers to a set of ten reference sequences with equal abundance; skew10 refers to ten reference sequences with two sequences 10 times as abundant as the rest. The plus sign suggests the clusters are validated with Meshclust.

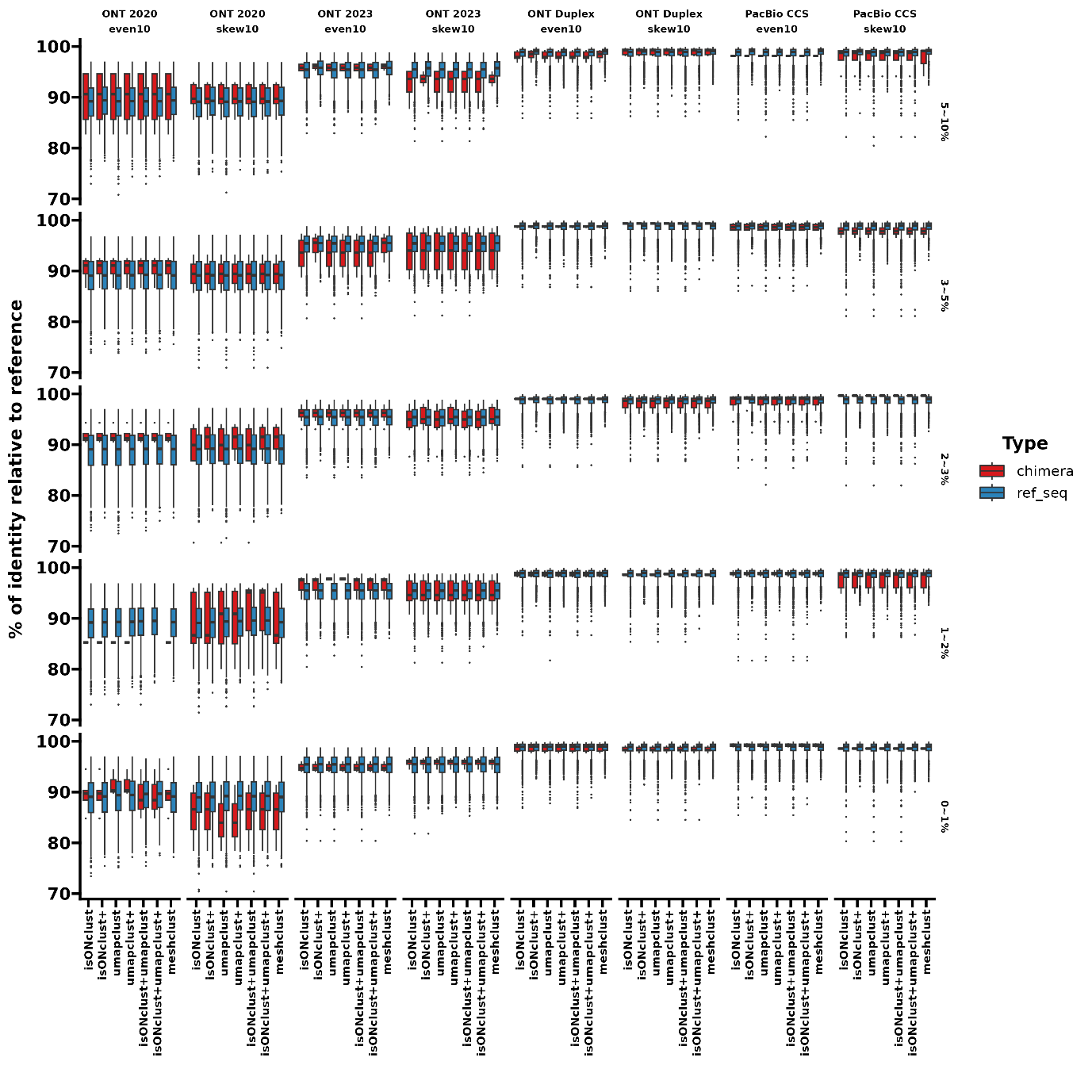

Supplementary Fig. 2. The sequence identity of clustered reads by various alignment-free approaches relative to the SILVA simulation reference sequences. The boxplot is colored by the simulation read types, based on the clustering results under the *in silico* sequencing depth of 200×. KmerCon, consensus calling on clustered sequences with UMAPclust and Meshclust; miniCon, consensus calling on clustered sequences refined by overlap check; isoCon, consensus calling of detected isoforms by IsoCon on the clustered sequences. The reference sequences are chosen from SILVA small subunit sequences with the divergence of 5-10%, 3-5%, 2-3% 1-2%, and 0-1%. Even10 refers to a set of ten reference sequences with equal abundance; skew10 refers to ten reference sequences with two sequences 10 times as abundant as the rest. The plus sign suggests the clusters are validated with Meshclust.

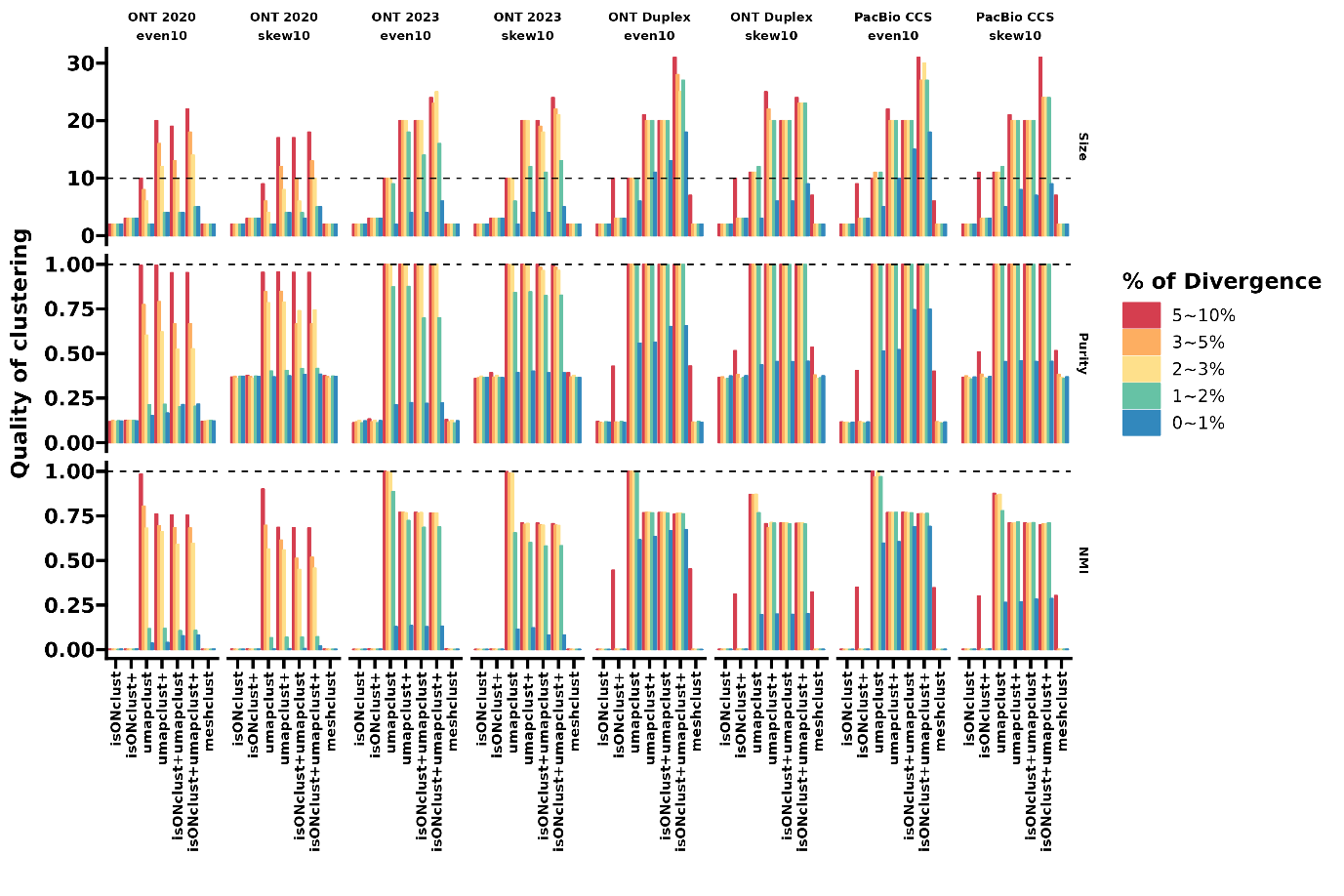

Supplementary Fig. 3. The clustering quality of various alignment-free approaches on the SILVA amplicons under the *in silico* sequencing depth of 200×. The clustering quality is assessed by size (the number of clusters), purity, and normalized mutual information (NMI) score. The black dashed lines suggest the theoretically optimal values, and the sequence orientation (strand) information is not considered in the clustering evaluation, resulting in the theoretical number of reference sequences of 10. The simulation template references are chosen from SILVA small subunit sequences with the divergence of 5-10%, 3-5%, 2-3% 1-2%, and 0-1%, and respective results are depicted by color. Even10 refers to a set of ten reference sequences with equal abundance; skew10 refers to ten reference sequences with two sequences 10 times as abundant as the rest. The plus sign suggests the clusters are validated with Meshclust.

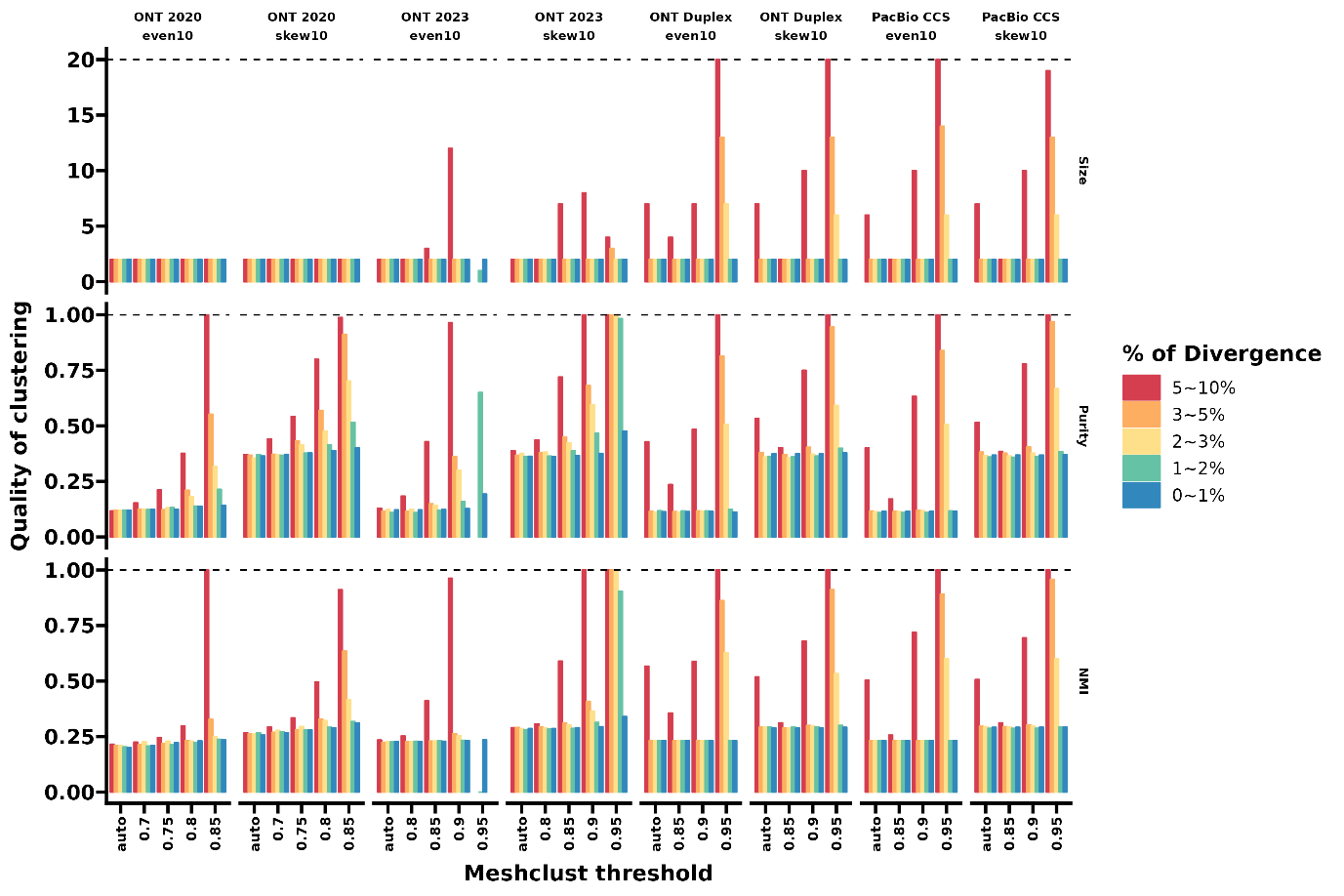

Supplementary Fig. 4. The effect of Meshclust thresholds (minimal clustering identities) on the clustering quality of the SILVA amplicons under the *in silico* sequencing depth of 200×. The clustering quality is assessed by size (the number of clusters), purity, and normalized mutual information (NMI) score. The black dashed lines suggest the theoretically optimal values. The simulation template references are chosen from SILVA small subunit sequences with the divergence of 5-10%, 3-5%, 2-3% 1-2%, and 0-1%, and respective results are depicted by color. Even10 refers to a set of ten reference sequences with equal abundance; skew10 refers to ten reference sequences with two sequences 10 times as abundant as the rest.

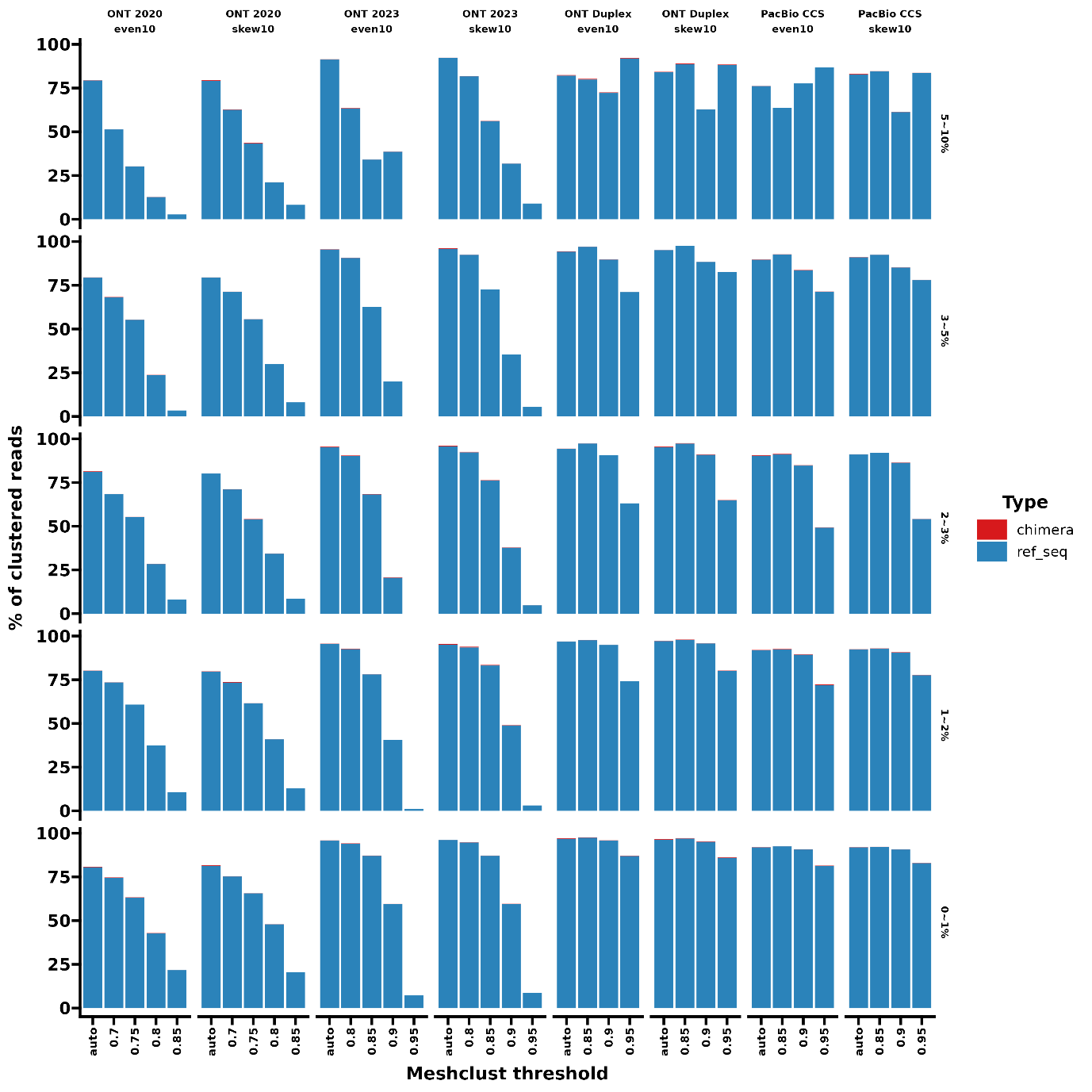

Supplementary Fig. 5. The portion of clustered reads with various Meshclust thresholds (minimal clustering identities) on the SILVA amplicons under the *in silico* sequencing depth of 200×. The portion is colored by the simulated read types. The reference sequences are chosen from SILVA small subunit sequences with the sequence divergence of 5-10%, 3-5%, 2-3% 1-2%, and 0-1%. Even10 refers to a set of ten reference sequences with equal abundance; skew10 refers to ten reference sequences with two sequences 10 times as abundant as the rest.

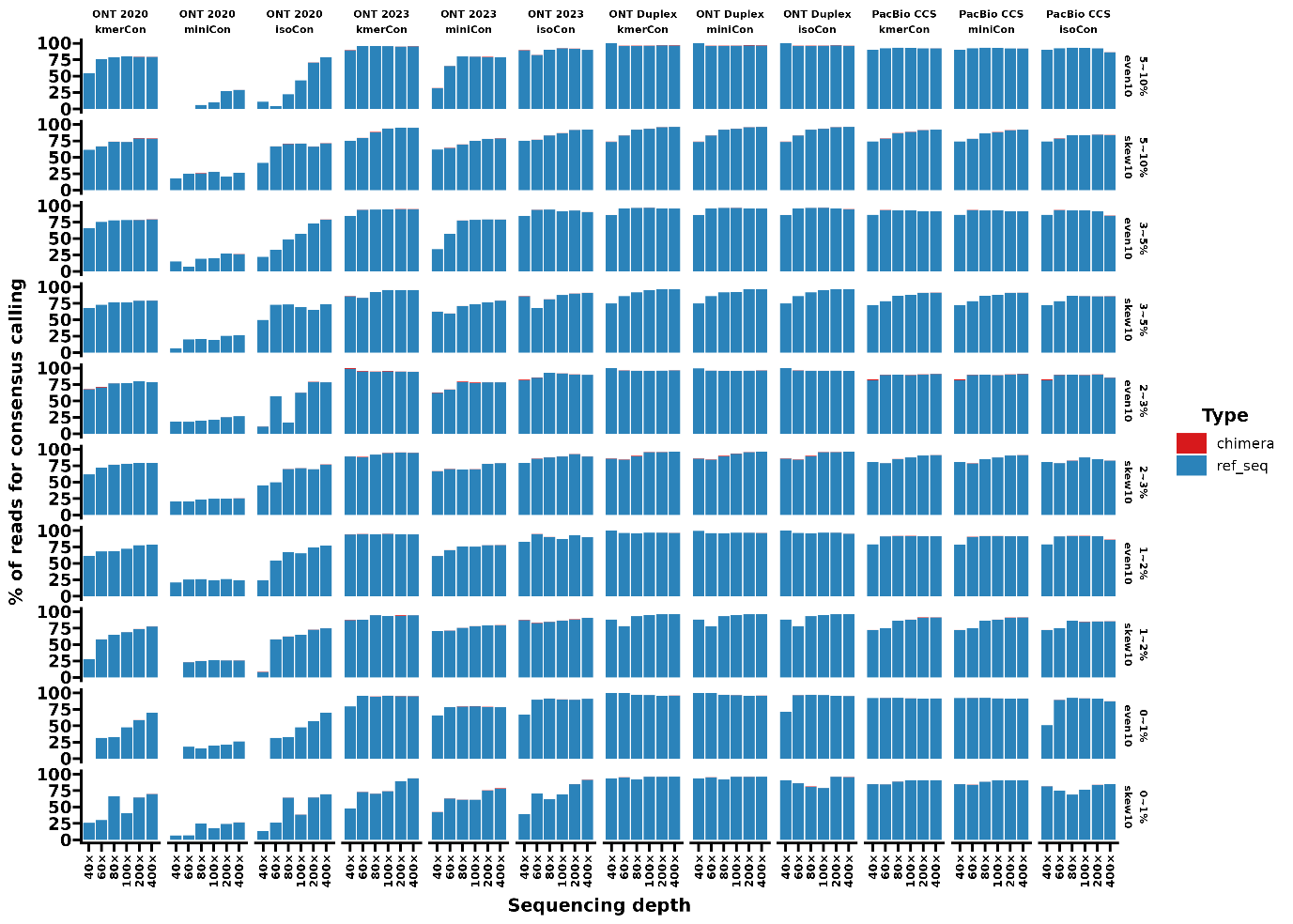

Supplementary Figure 6. The portion of utilized reads with the three consensus-calling approaches on SILVA amplicons under the *in silico* sequencing depth from 40× to 400×. The portion is colored by the simulated read types. The reference sequences are chosen from SILVA small subunit sequences with the divergence of 5-10%, 3-5%, 2-3% 1-2%, and 0-1%. KmerCon, consensus calling on clustered sequences with UMAPclust and Meshclust; miniCon, consensus calling on clustered sequences refined by overlap check; isoCon, consensus calling of detected isoforms by IsoCon on the clustered sequences. Even10 refers to a set of ten reference sequences with equal abundance; skew10 refers to ten reference sequences with two sequences 10 times as abundant as the rest.

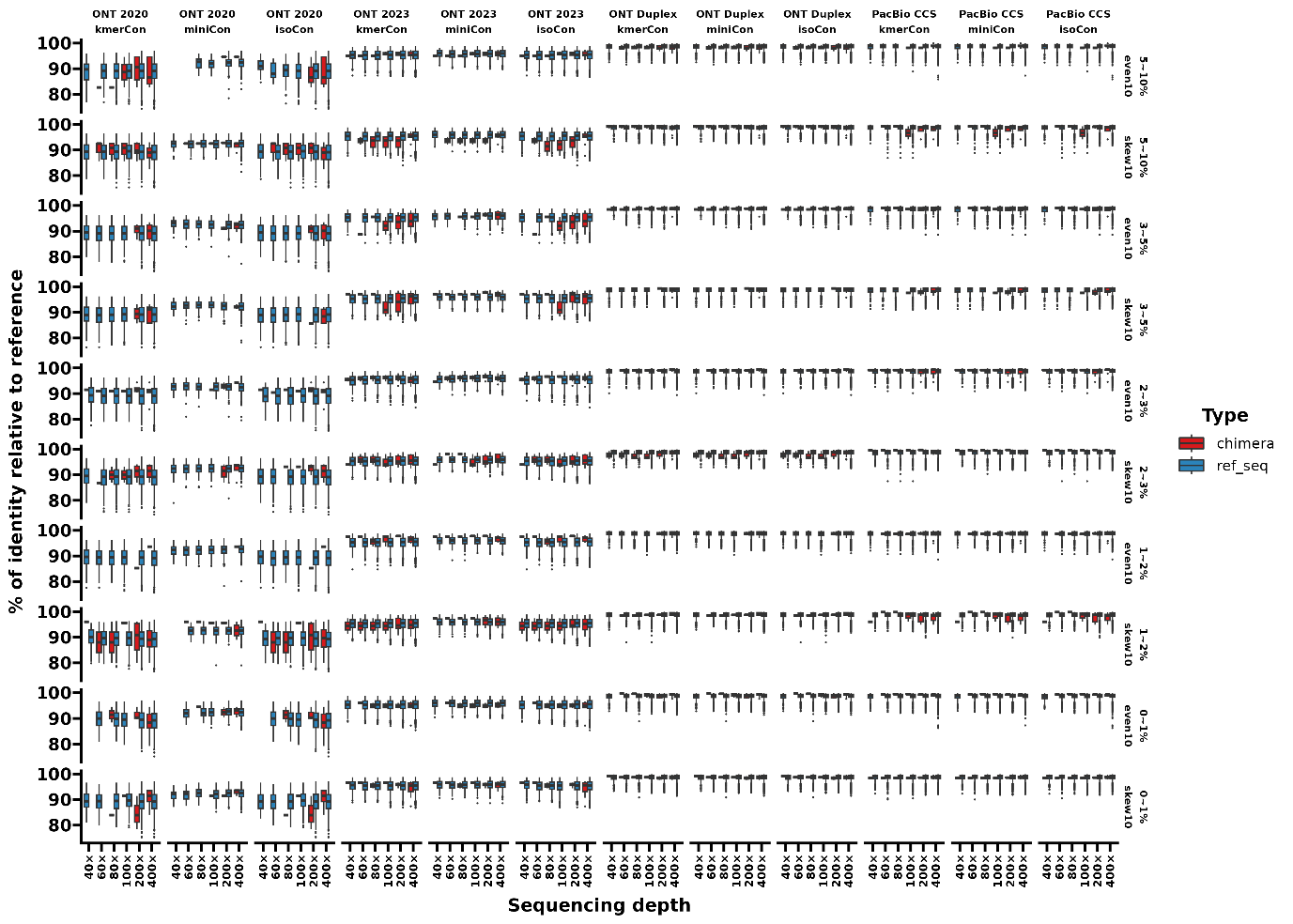

Supplementary Figure 7. The sequence identity of picked reads by the three consensus calling approaches relative to the SILVA simulation reference sequences. The boxplot is colored by the simulation read types, based on the results under the *in silico* sequencing depth

from 40× to 400×. The reference sequences are chosen from SILVA small subunit sequences with the divergence of 5-10%, 3-5%, 2-3% 1-2%, and 0-1%. KmerCon, consensus calling on clustered sequences with UMAPclust and Meshclust; miniCon, consensus calling on clustered sequences refined by overlap check; isoCon, consensus calling of detected isoforms by IsoCon on the clustered sequences. Even10 refers to a set of ten reference sequences with equal abundance; skew10 refers to ten reference sequences with two sequences 10 times as abundant as the rest.

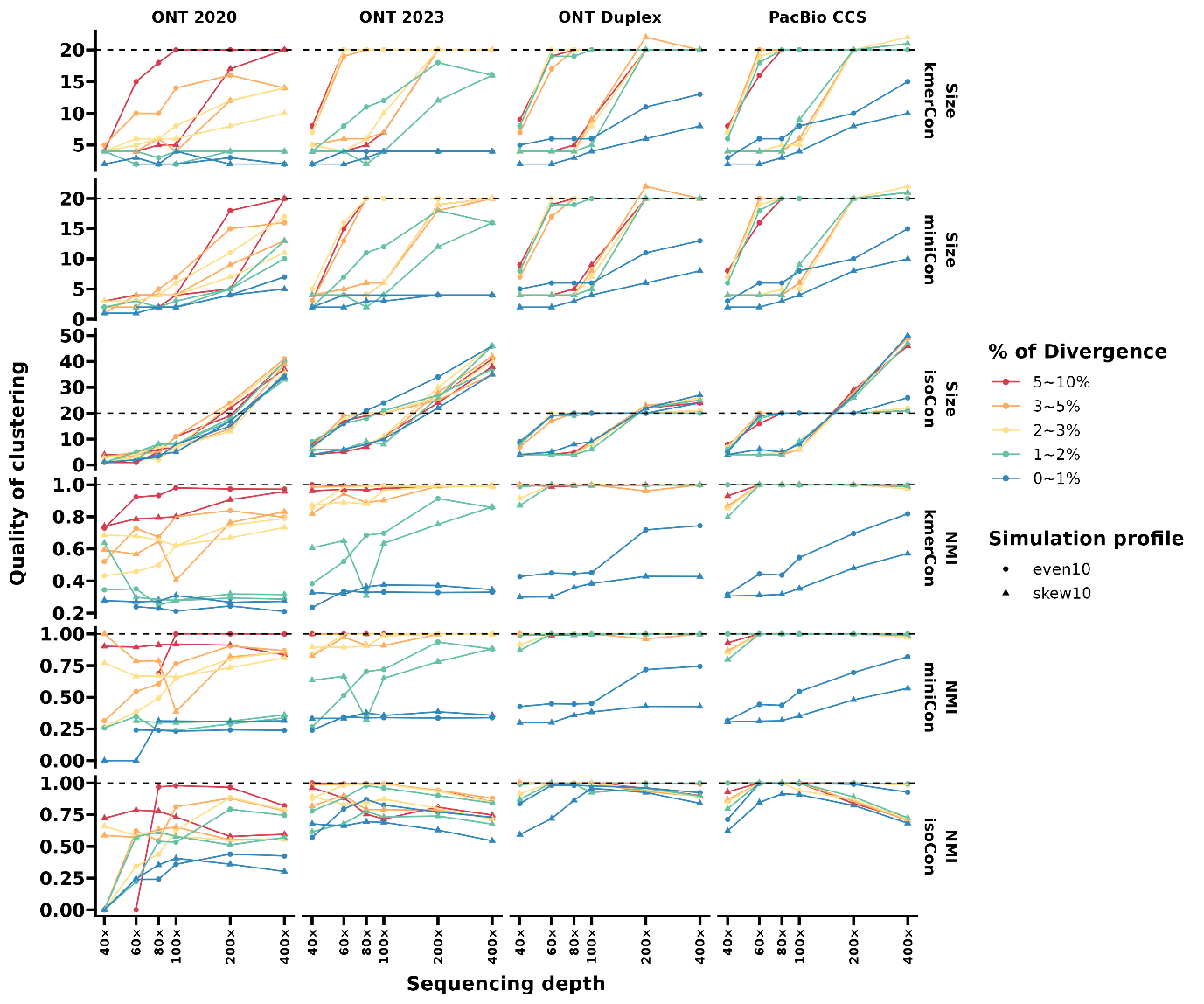

Supplementary Fig. 8. The number (a) and the normalized mutual information (NMI) score (b) of the validated clusters from the three consensus calling approaches on the SILVA amplicons under the *in silico* sequencing depth from 40× to 400×. The black dashed lines suggest the theoretically optimal values, and the sequence orientation (strand) information is considered in the clustering evaluation, resulting in the theoretical number of reference sequences of 20. The simulation template references are chosen from SILVA small subunit sequences with the divergence of 5-10%, 3-5%, 2-3% 1-2%, and 0-1%, and respective results are depicted by color. The point shapes indicate the simulation scenarios of reference sequences in even (dots) and skewed (triangles) composition. KmerCon, consensus calling on clustered sequences with UMAPclust and Meshclust; miniCon, consensus calling on clustered sequences refined by overlap check; isoCon, consensus calling of detected isoforms by IsoCon on the clustered sequences. Even10 refers to a set of ten reference sequences with equal abundance; skew10 refers to ten reference sequences with two sequences 10 times as abundant as the rest.

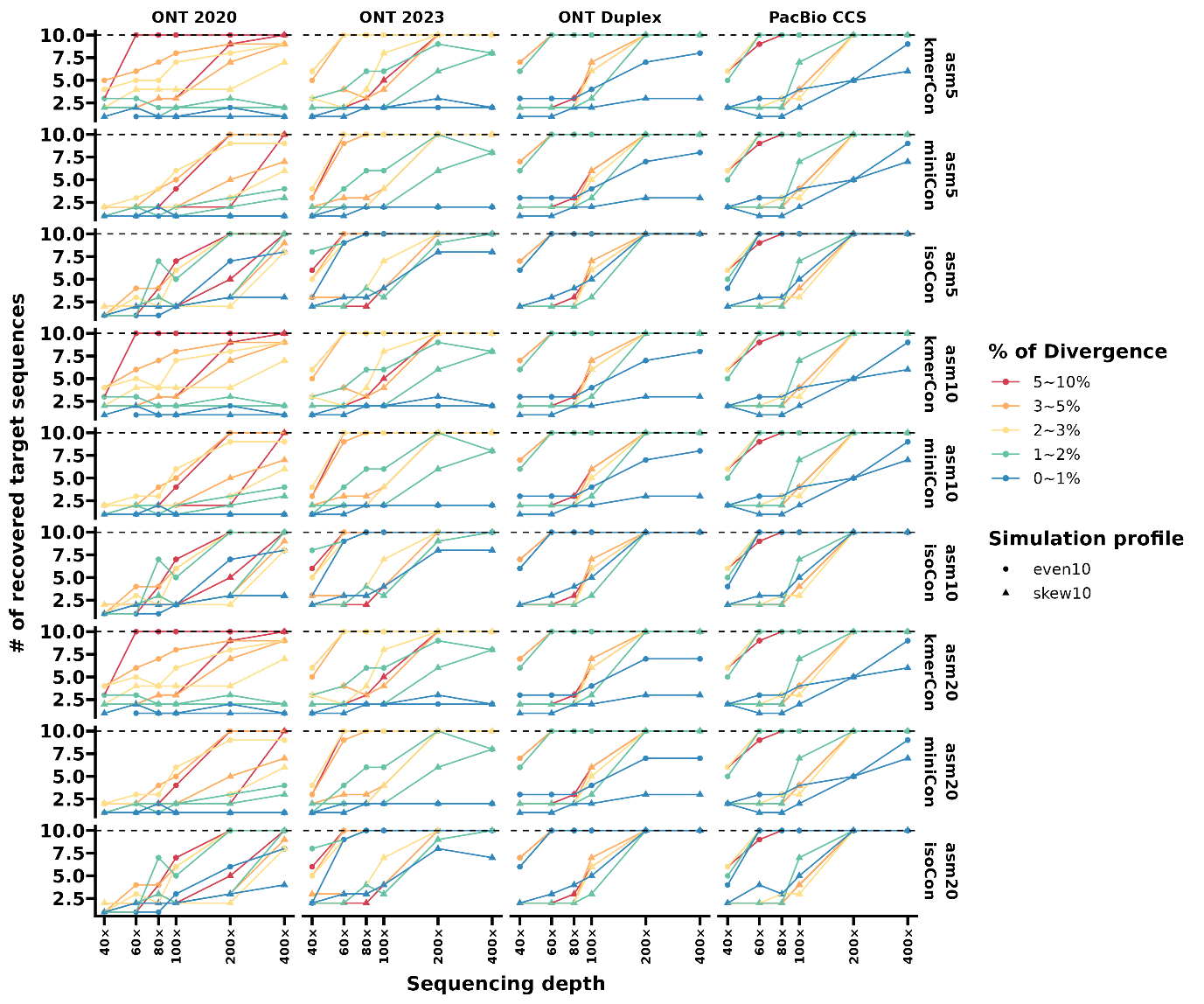

Supplementary Fig. 9. The number of retrieved reference sequences using the three consensus calling approaches on the SILVA amplicons under the *in silico* sequencing depth from 40× to 400×. The black dashed lines suggest the theoretically optimal values. The simulation template references are chosen from SILVA small subunit sequences with the divergence of 5-10%, 3-5%, 2-3% 1-2%, and 0-1%, and respective results are depicted by color. The point shapes indicate the simulation scenarios of reference sequences in even (dots) and skewed (triangles) composition. KmerCon, consensus calling on clustered sequences with UMAPclust and Meshclust; miniCon, consensus calling on clustered sequences refined by overlap check; isoCon, consensus calling of detected isoforms by IsoCon on the clustered sequences. Even10 refers to a set of ten reference sequences with equal abundance; skew10 refers to ten reference sequences with two sequences 10 times as abundant as the rest.

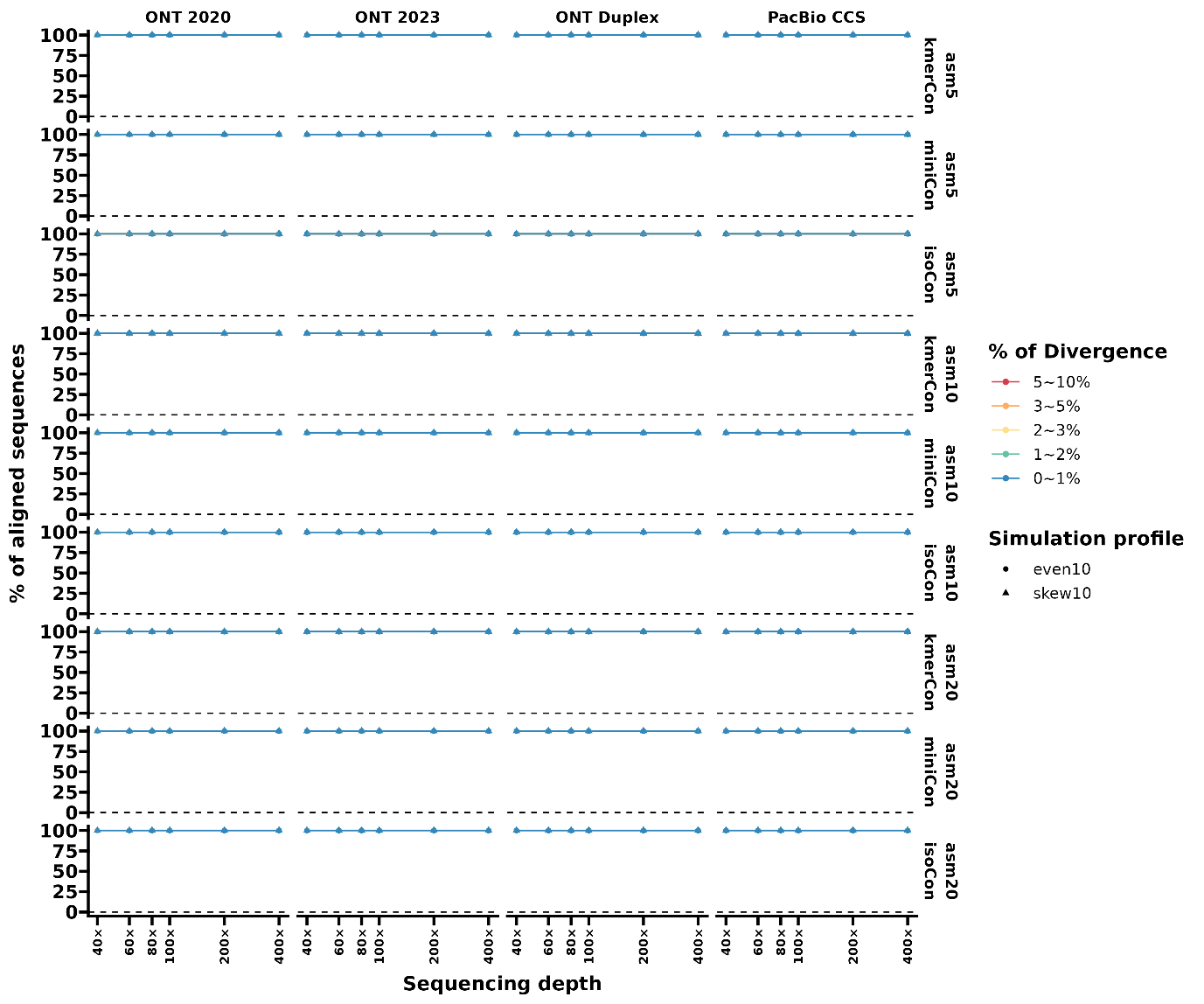

Supplementary Figure 10. The percentage of the consensus sequences aligned to respective the SILVA simulation reference sequences. The plot is based on the results under the *in silico* sequencing depth from 40× to 400×. The black dashed lines suggest the theoretically optimal values. The simulation template references are chosen from SILVA small subunit sequences with the divergence of 5-10%, 3-5%, 2-3% 1-2%, and 0-1%, and respective results are depicted by color. The point shapes indicate the simulation scenarios of reference sequences in even (dots) and skewed (triangles) composition. KmerCon, consensus calling on clustered sequences with UMAPclust and Meshclust; miniCon, consensus calling on clustered sequences refined by overlap check; isoCon, consensus calling of detected isoforms by IsoCon on the clustered sequences. Even10 refers to a set of ten reference sequences with equal abundance; skew10 refers to ten reference sequences with two sequences 10 times as abundant as the rest.

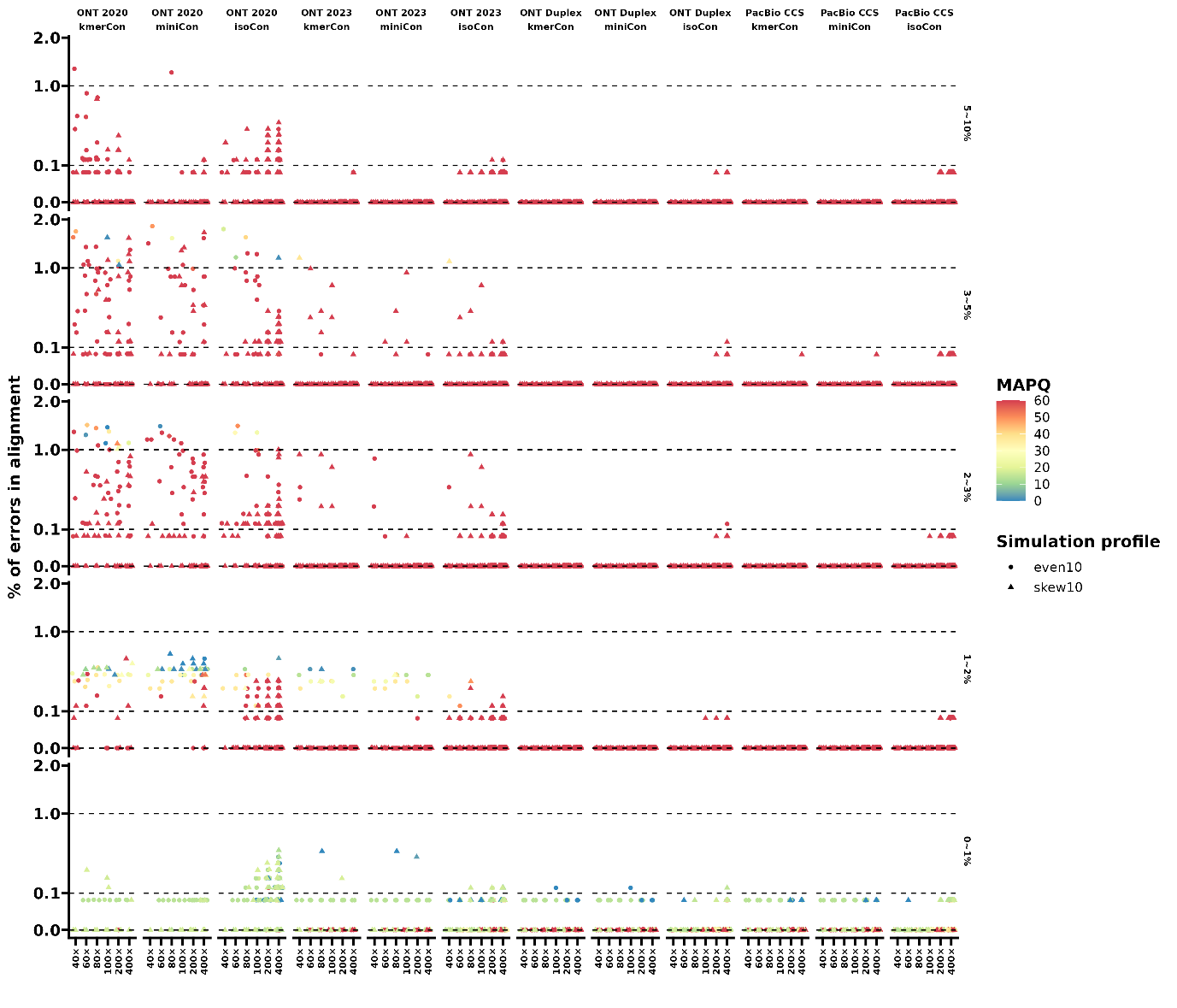

Supplementary Figure 11. The percentage of errors in alignment between the corrected sequences and the respective SILVA reference sequences under the *in silico* sequencing depth from 40× to 400×. Square root scaling is employed to improve the visualization of the error rate. The point shapes indicate the simulation scenarios of reference sequences in even (dots) and skewed (triangles) composition, and color depth suggests the mapping quality score (MAPQ) from minimap2. KmerCon, consensus calling on clustered sequences with UMAPclust and Meshclust; miniCon, consensus calling on clustered sequences refined by overlap check; isoCon, consensus calling of detected isoforms by IsoCon on the clustered sequences. Even10 refers to a set of ten reference sequences with equal abundance; skew10 refers to ten reference sequences with two sequences 10 times as abundant as the rest.

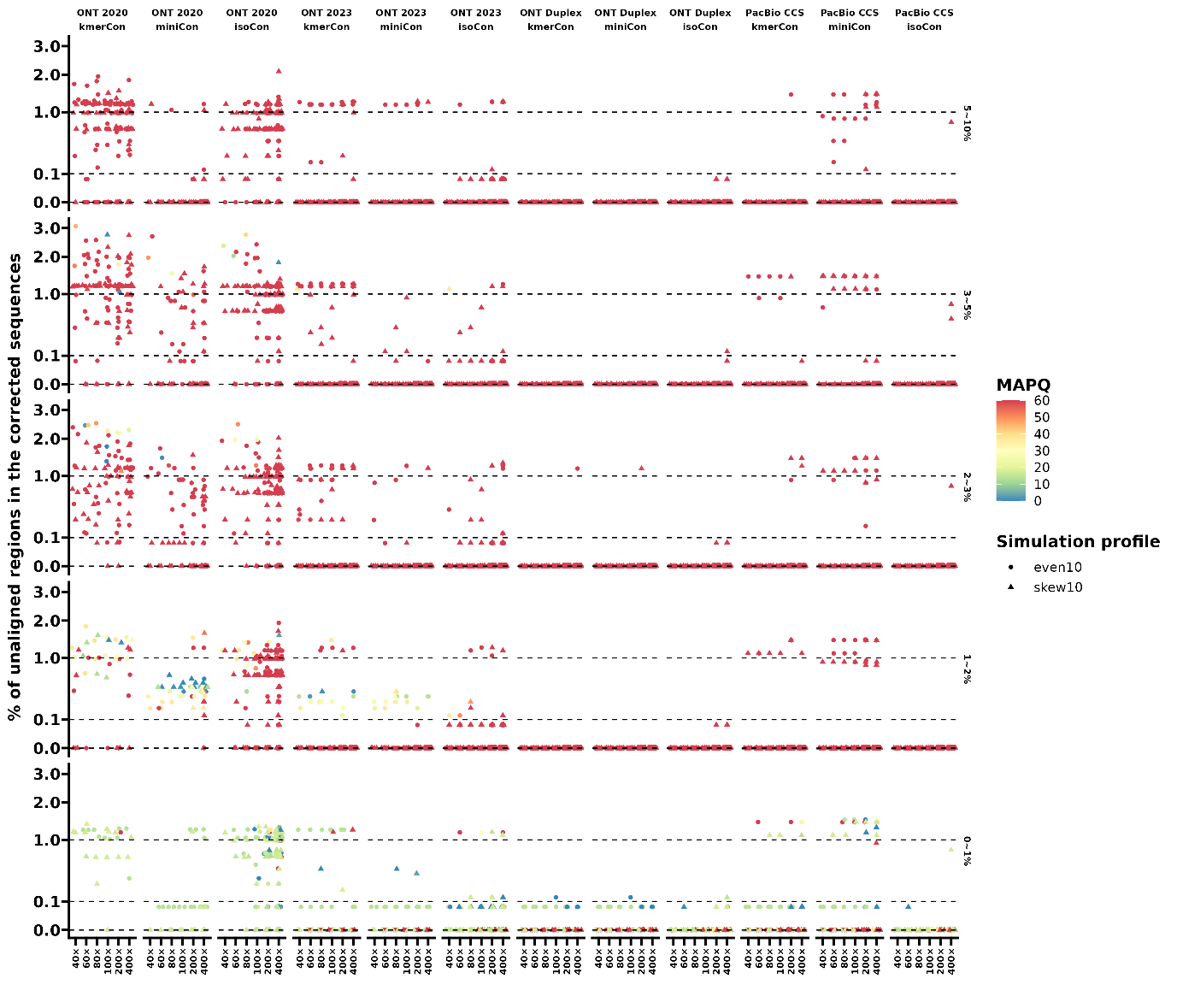

Supplementary Figure 12. The percentage of unaligned regions in the corrected sequence to the respective reference sequences under the *in silico* sequencing depth from 40× to 400×. Square root scaling is employed to improve the visualization of the unalignment percentage. The point shapes indicate the simulation scenarios of reference sequences in even (dots) and skewed (triangles) composition, and color depth suggests the mapping quality score (MAPQ) from minimap2. KmerCon, consensus calling on clustered sequences with UMAPclust and Meshclust; miniCon, consensus calling on clustered sequences refined by overlap check; isoCon, consensus calling of detected isoforms by IsoCon on the clustered sequences. Even10 refers to a set of ten reference sequences with equal abundance; skew10 refers to ten reference sequences with two sequences 10 times as abundant as the rest.

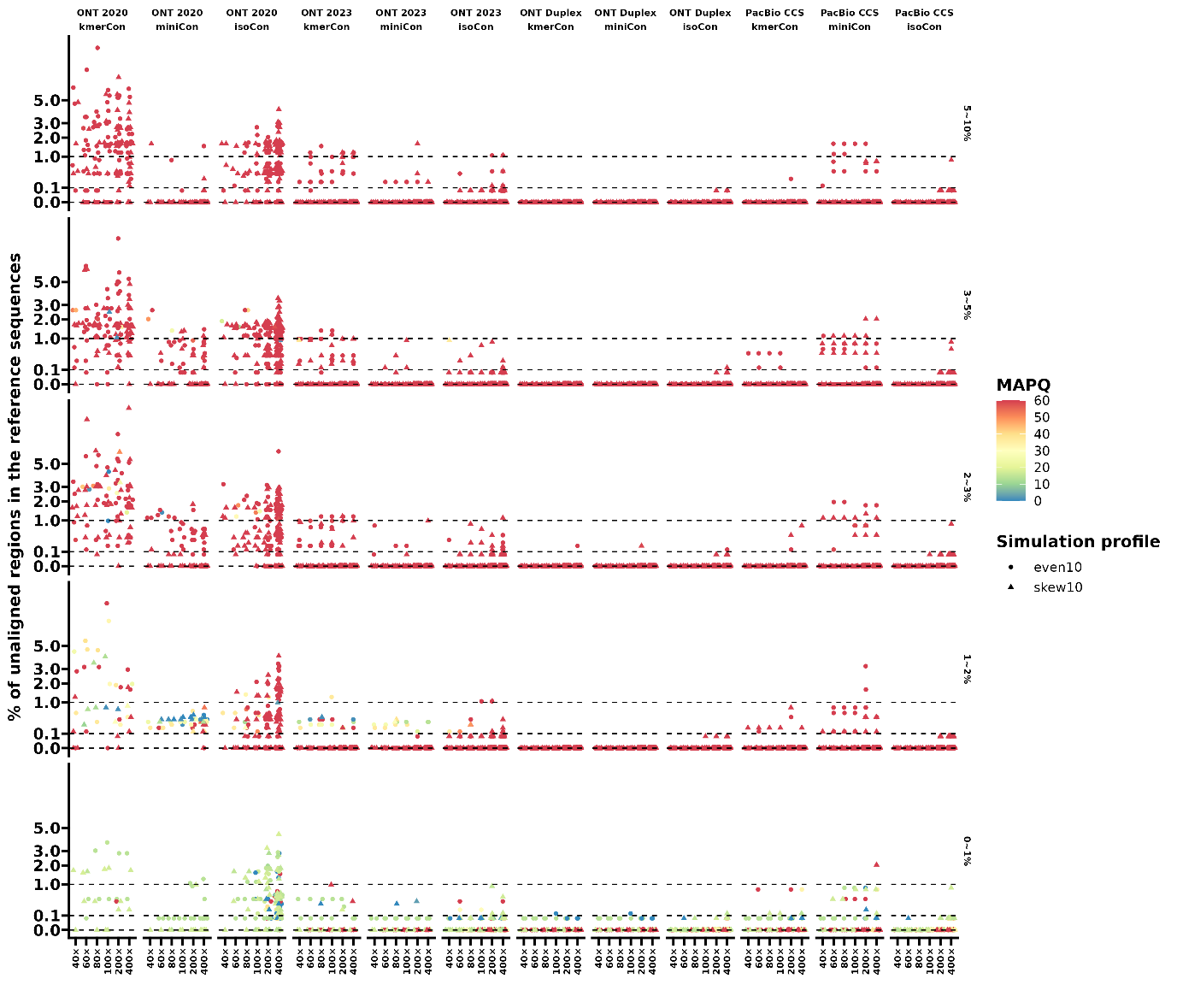

Supplementary Figure 13. The percentage of unaligned regions in the reference sequences under the *in silico* sequencing depth from 40× to 400×. Square root scaling is employed to improve the visualization of the unalignment percentage. The point shapes indicate the simulation scenarios of reference sequences in even (dots) and skewed (triangles) composition, and color depth suggests the mapping quality score (MAPQ) from minimap2. KmerCon, consensus calling on clustered sequences with UMAPclust and Meshclust; miniCon, consensus calling on clustered sequences refined by overlap check; isoCon, consensus calling of detected isoforms by IsoCon on the clustered sequences. Even10 refers to a set of ten reference sequences with equal abundance; skew10 refers to ten reference sequences with two sequences 10 times as abundant as the rest.

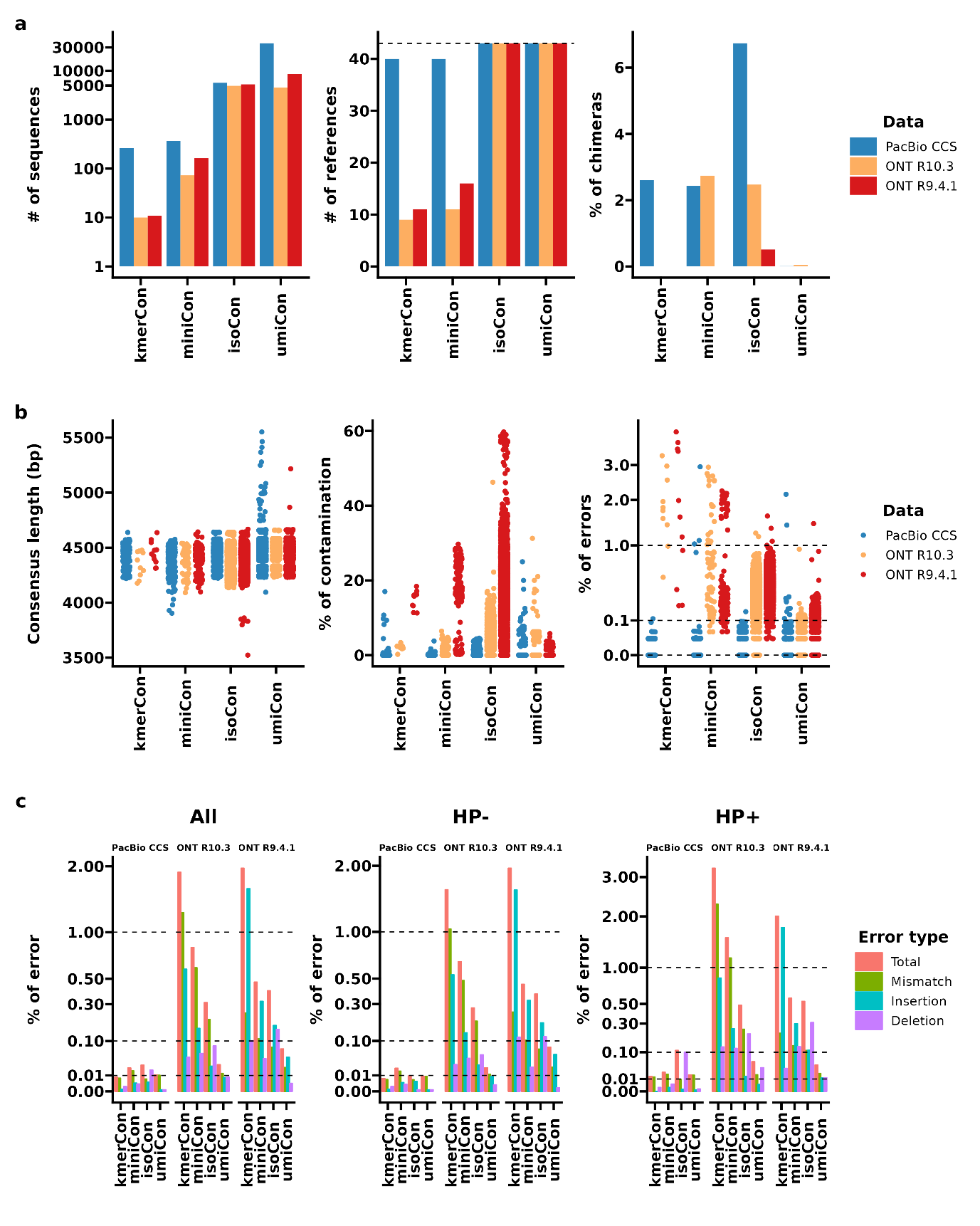

Supplementary Figure 14. Statistics of various amplicon consensus sequences retrieved from the subsampled 1,000,000 reads tagged with unique molecular identifiers (UMIs). a, The number of consensus sequences and the identified reference sequences, and the chimera rate; b. The sequence length, the percentage of reads aligned to a reference sequence other than the primary class (contamination), and the error rate of each consensus sequence; c, The average error rate for mismatches, insertions, deletions, and total errors of retrieved consensus sequences, split by whether the error occurred inside (hp+) or outside (hp-) a homopolymer (hp) region. The consensus sequences are quality-controlled with PCR artifacts removed and kept with a coverage cutoff according to the consensus modes and sequencing platforms. The coverage cutoffs of kmerCon and miniCon are 400× and 300×, while the cutoffs are 20×, 20×, and 25× for isoCon and 3×, 15×, and 25× for umiCon on the sequencing data from PacBio CCS, ONT R10.3 and ONT R9.4.1, respectively. To enhance visualization, we applied a base-10 logarithmic scaling to the number of consensus sequences in sub-figure (a) and employed square root scaling for the error rates in sub-figures (b) and (c). The three datasets are colored by the sequencing platforms in sub-figure (a) and (b), and the bar colors in sub-figure (c) display different error categories. KmerCon, consensus calling on clustered sequences with UMAPclust and Meshclust; miniCon, consensus calling on clustered sequences refined by overlap check; isoCon, consensus calling of detected isoforms by IsoCon on the clustered sequences; umiCon, consensus calling on UMI bins.

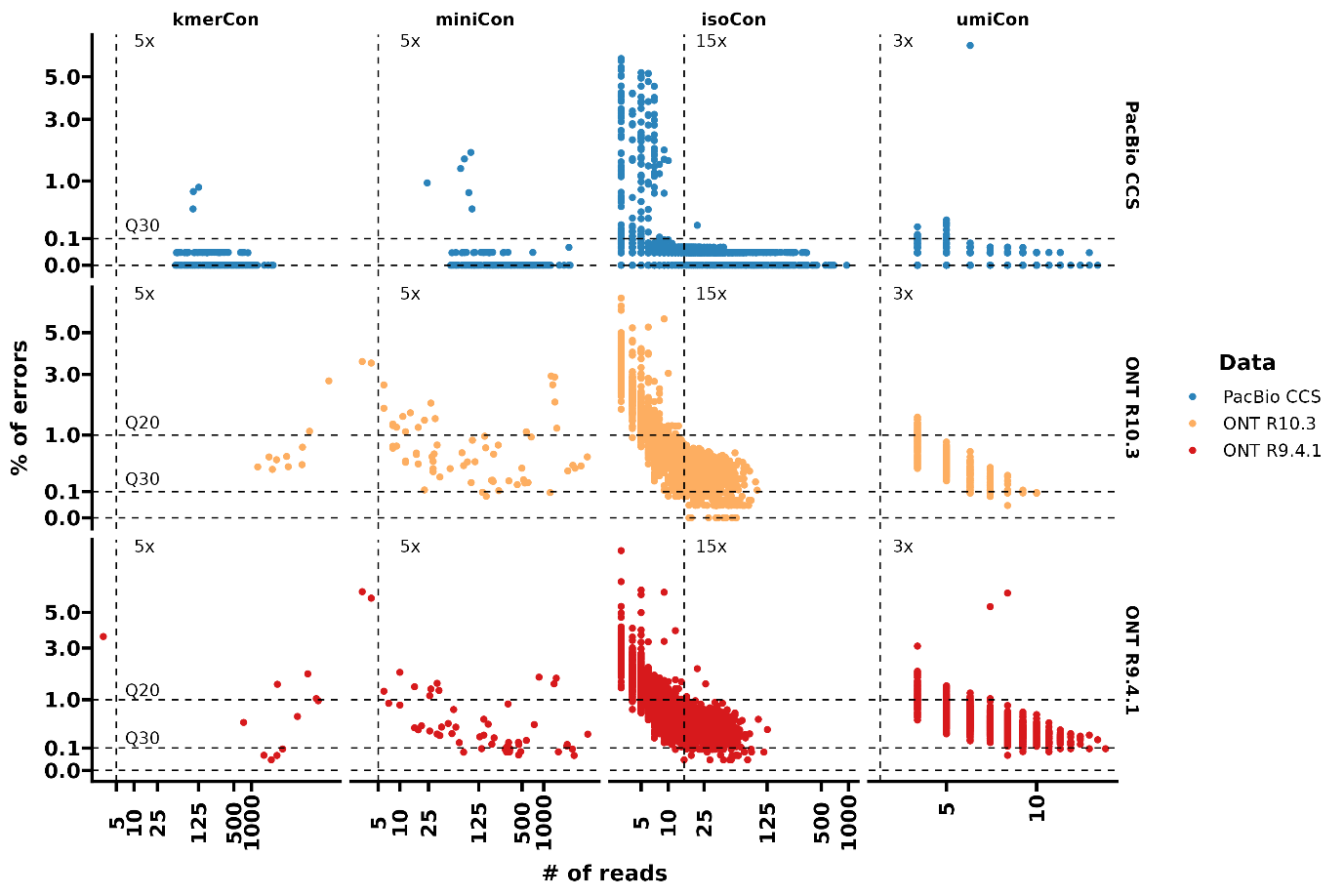

Supplementary Figure 15. The error rate and read coverage of each amplicon consensus sequence retrieved from the subsampled 100,000 reads tagged with unique molecular identifiers (UMIs). Square root scaling is employed to improve the visualization of the error rate. The three datasets are colored by the sequencing platforms, the dotted vertical lines suggest the read coverage cutoff for respective consensus sequence sets. KmerCon, consensus calling on clustered sequences with UMAPclust and Meshclust; miniCon, consensus calling on clustered sequences refined by overlap check; isoCon, consensus calling of detected isoforms by IsoCon on the clustered sequences; umiCon, consensus calling on UMI bins.

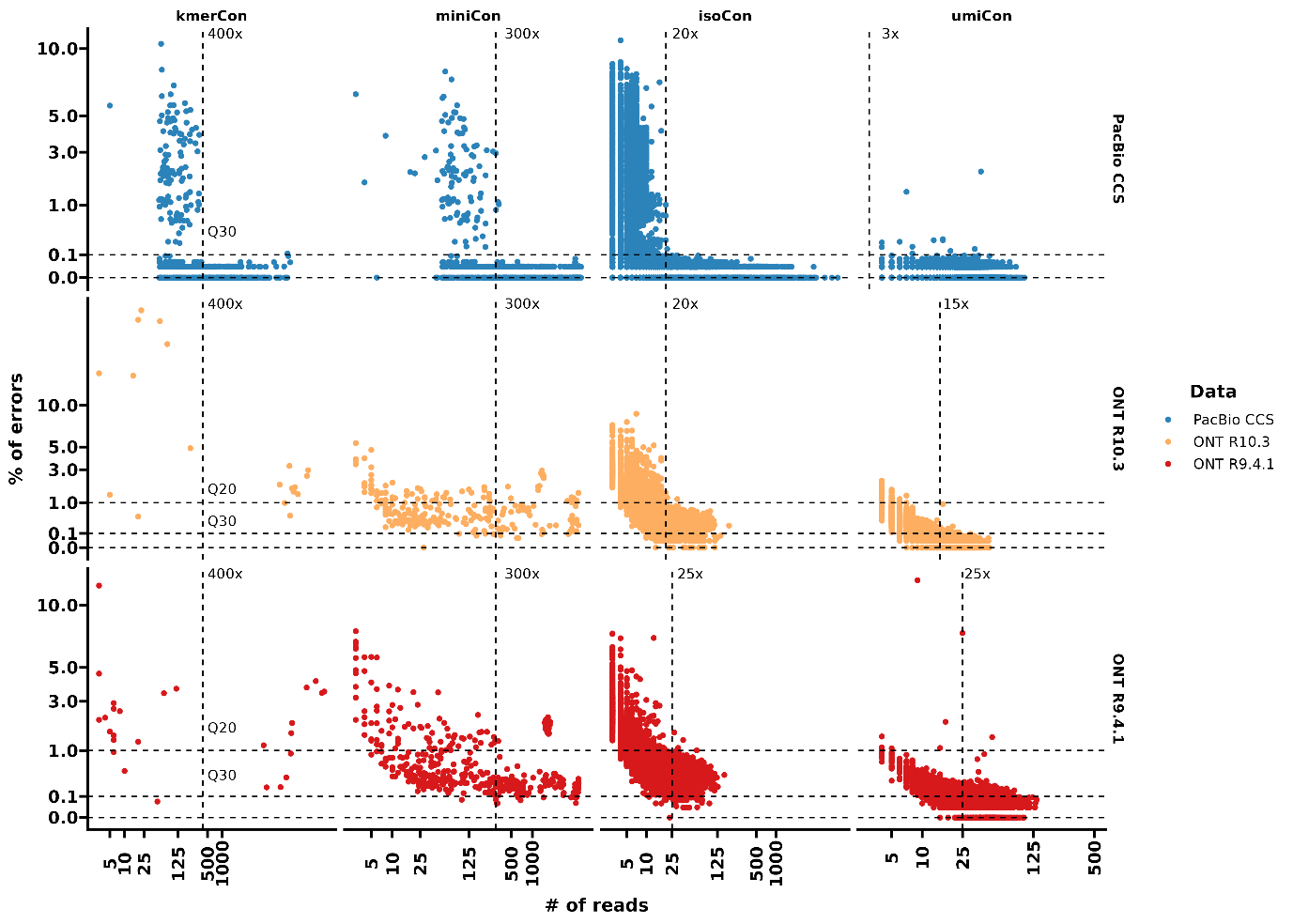

Supplementary Figure 16. The error rate and read coverage of each amplicon consensus sequence retrieved from the subsampled 1,000,000 reads tagged with unique molecular identifiers (UMIs). Square root scaling is employed to improve the visualization of the error rate. The three datasets are colored by the sequencing platforms, the dotted vertical lines suggest the read coverage cutoff for respective consensus sequence sets. KmerCon, consensus calling on clustered sequences with UMAPclust and Meshclust; miniCon, consensus calling on clustered sequences refined by overlap check; isoCon, consensus calling of detected isoforms by IsoCon on the clustered sequences; umiCon, consensus calling on UMI bins.

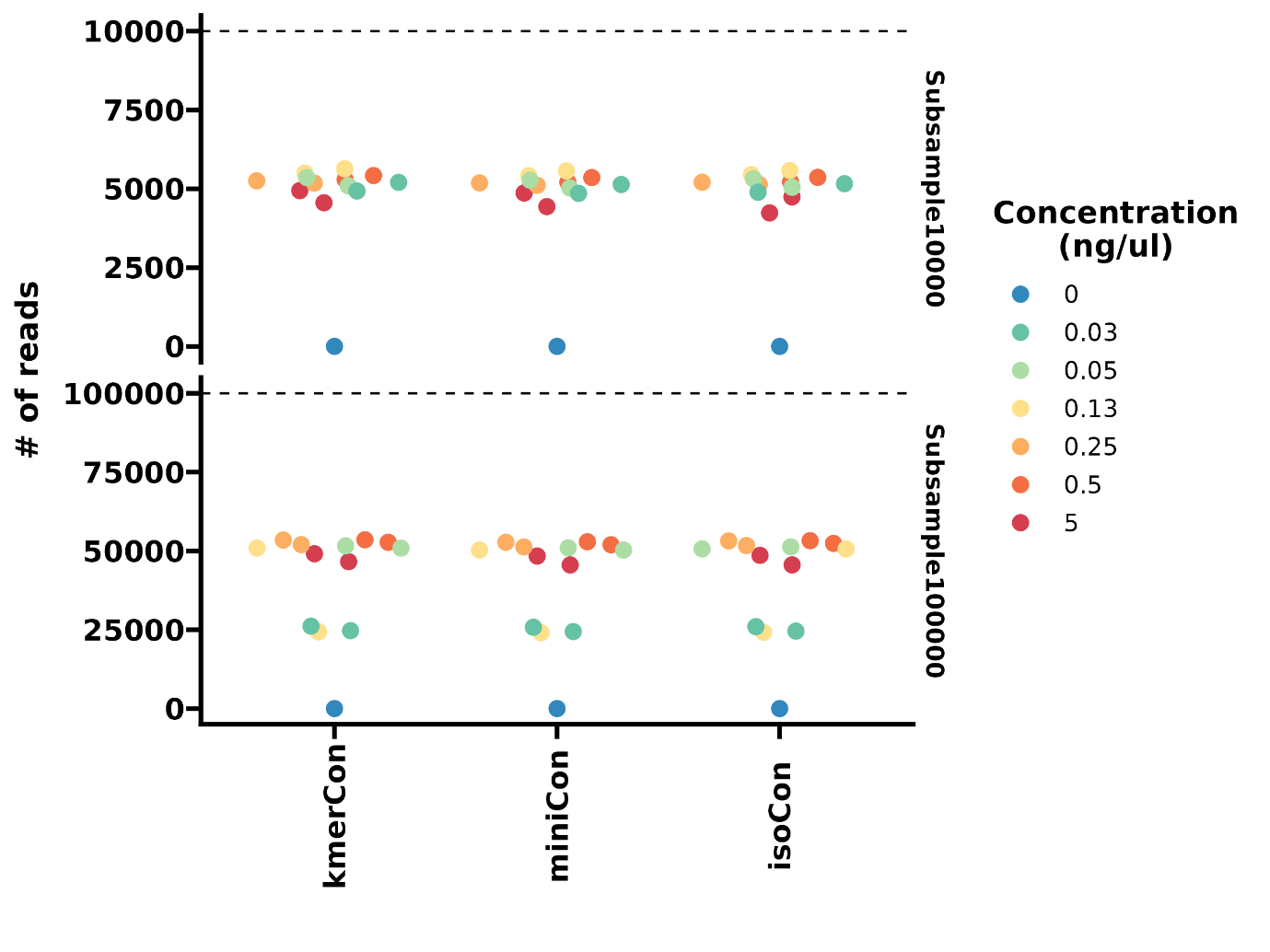

Supplementary Figure 17. The number of utilized reads by three OTU-picking approaches on mock serials with subsampled reads of 10,000 and 100,000 per sample. The OTUs are colored by the mock concentration in the dilution series. KmerCon, community profile based on clustered sequences with UMAPclust and Meshclust; miniCon, community profile based on clustered sequences refined with overlap check; isoCon, community profile based on detected isoforms by IsoCon on the clustered sequences.

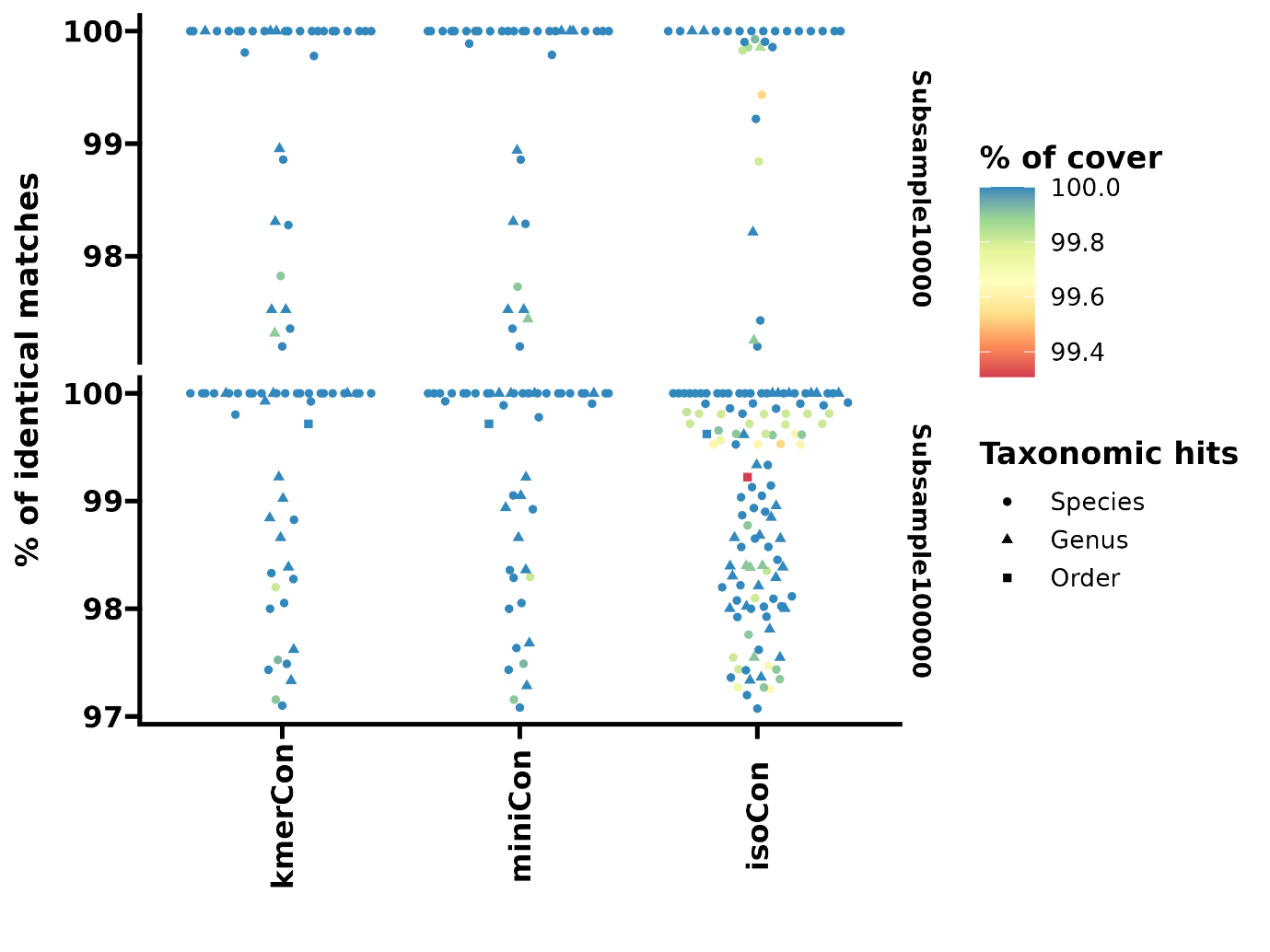

Supplementary Figure 18. The BLAST alignment identity of OTU sequences against the SILVA small subunit rRNA sequences. The point is colored by the cover percentage in BLAST alignment and shaped based on the lowest level of consensus taxonomic hits. KmerCon, community profile based on clustered sequences with UMAPclust and Meshclust; miniCon, community profile based on clustered sequences refined with overlap check; isoCon, community profile based on detected isoforms by IsoCon on the clustered sequences.

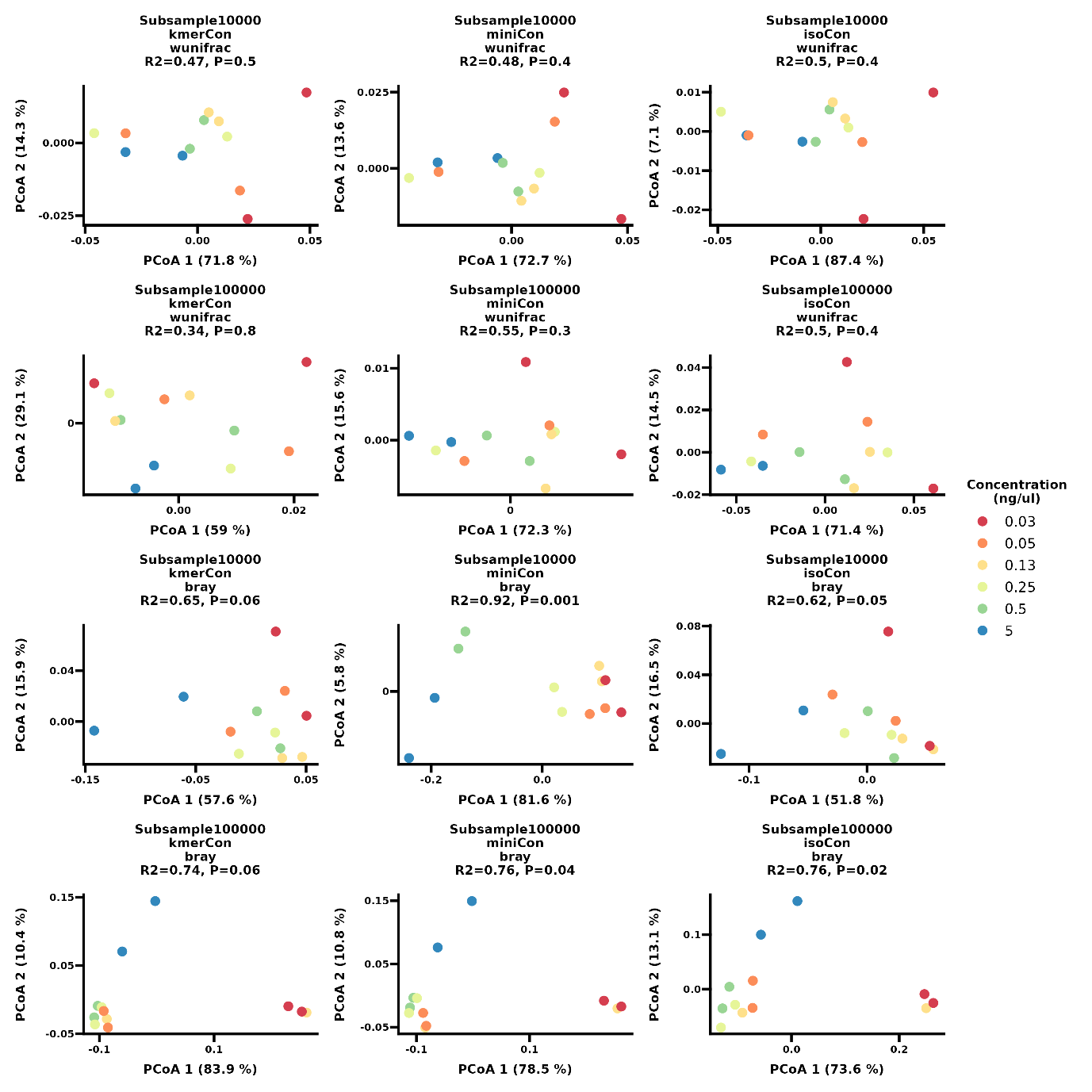

Supplementary Figure 19. Principal coordinate analysis (PCoA) plots of weighted UniFrac and Bray Curtis dissimilarity using the three OTU-picking methods on near full-length 16S rRNA amplicons of diluted mock serials at a subsampling depth of 10,000 and 100,000 reads per sample. The points are colored in accordance with respective mock concentrations in the dilution series. KmerCon, community profile based on clustered sequences with UMAPclust and Meshclust; miniCon, community profile based on clustered sequences refined with overlap check; isoCon, community profile based on detected isoforms by IsoCon on the clustered sequences.

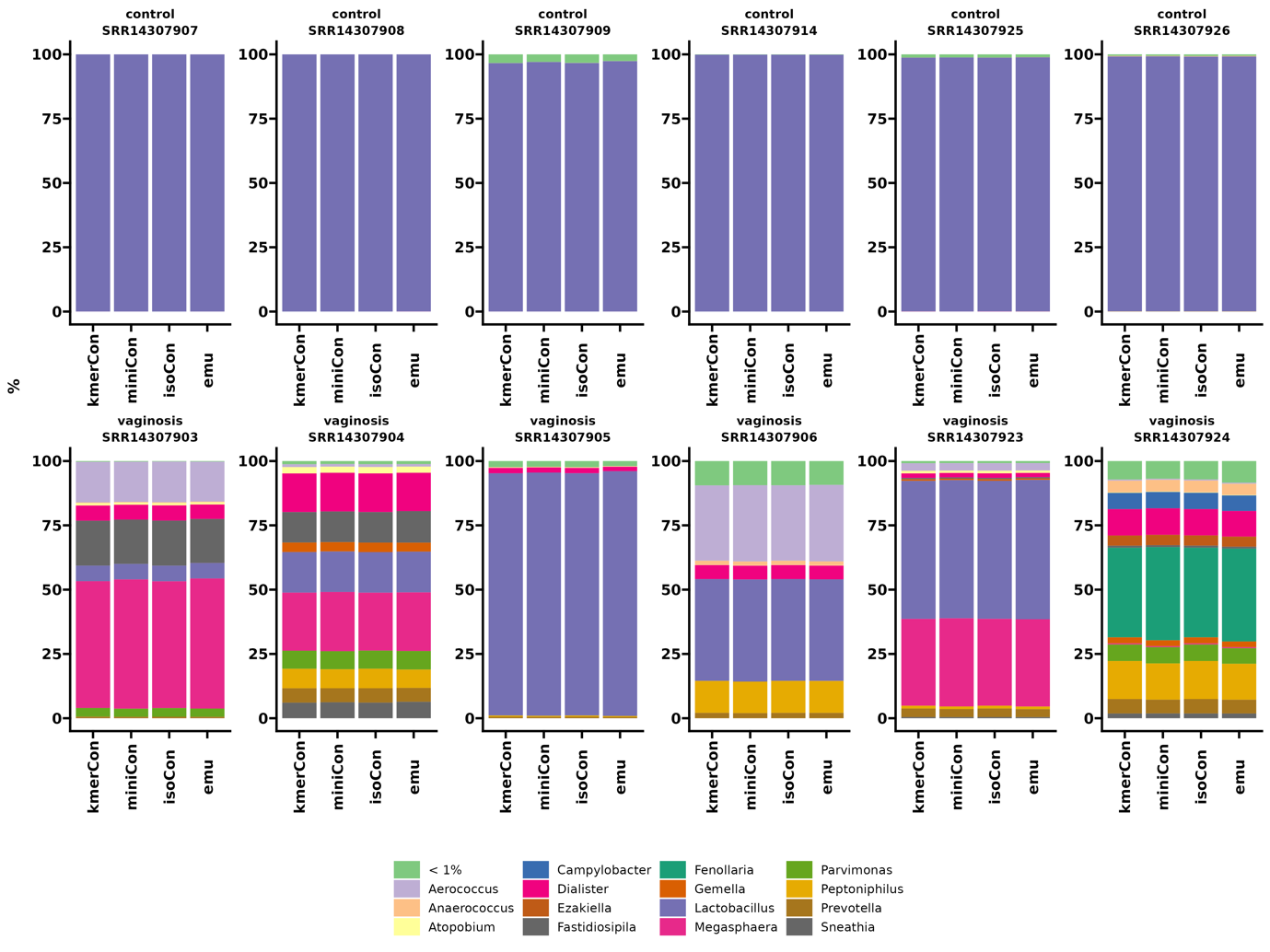

Supplementary Figure 20. Various genus-level profiles of human vaginal microbiomes with LACA and Emu. Taxonomic hits with minimum mean relative abundance of 1% are shown. Emu, read classification profile with Emu; kmerCon, community profile based on clustered sequences with UMAPclust and Meshclust; miniCon, community profile based on clustered sequences refined with overlap check; isoCon, community profile based on detected isoforms by IsoCon on the clustered sequences.

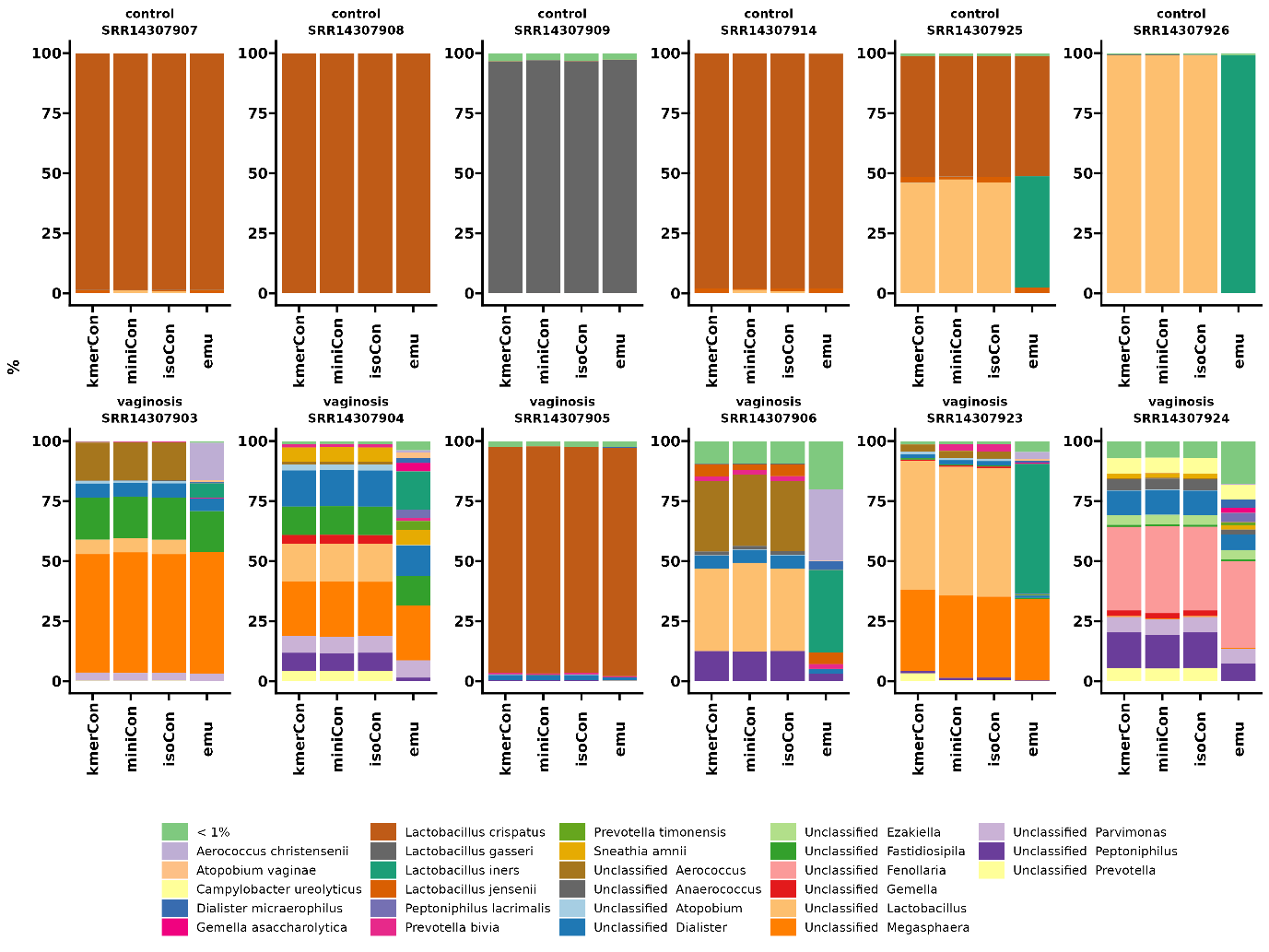

Supplementary Figure 21. Various species-level profiles of human vaginal microbiomes with LACA and Emu. Taxonomic hits with minimum mean relative abundance of 1% are shown. Emu, read classification profile with Emu; kmerCon, community profile based on clustered sequences with UMAPclust and Meshclust; miniCon, community profile based on clustered sequences refined with overlap check; isoCon, community profile based on detected isoforms by IsoCon on the clustered sequences.

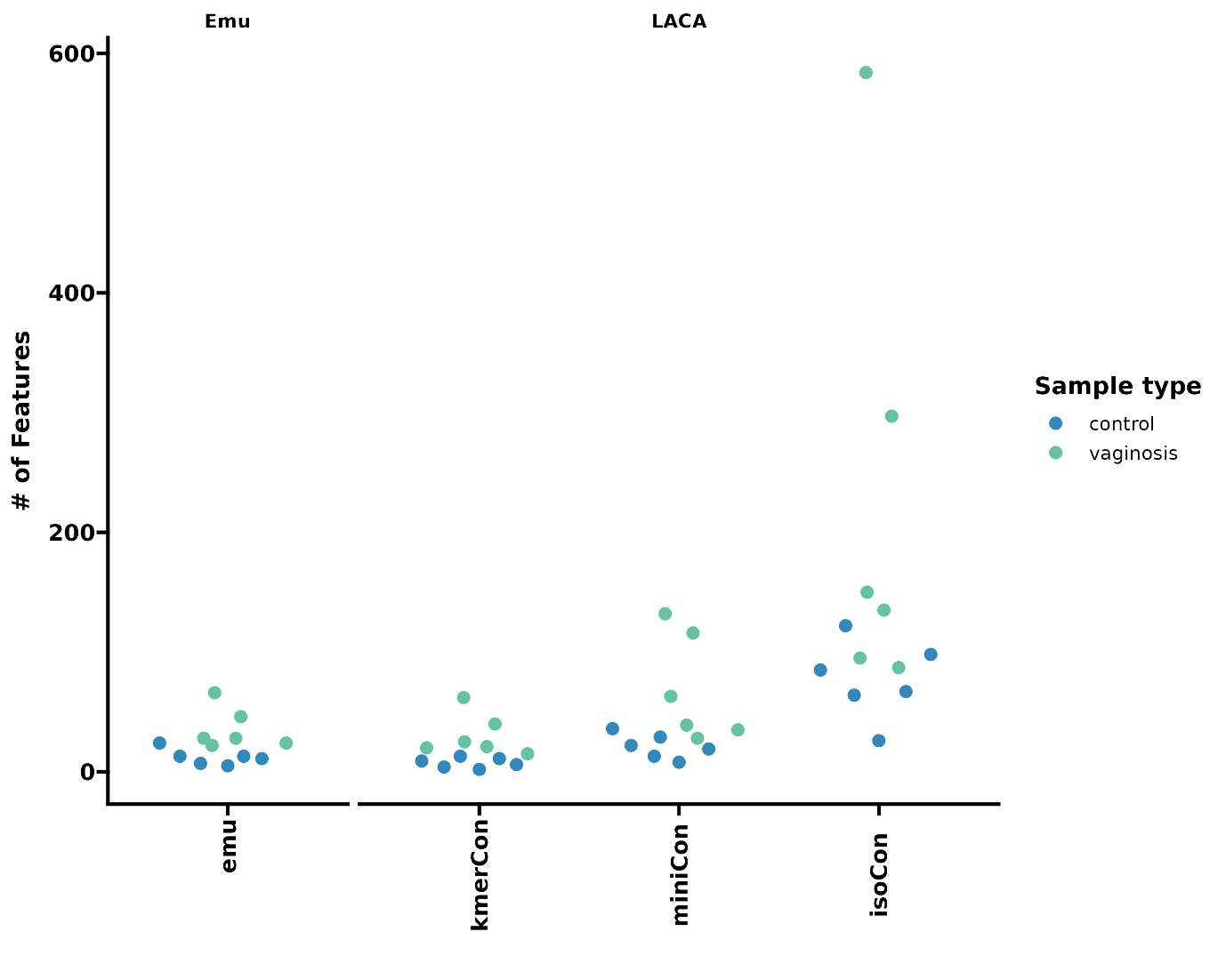

Supplementary Figure 22. The number of identified features by various profiling approaches on the human vaginal samples. The samples are shown in dots and colored by their phenotypes. Emu, read classification profile with Emu; kmerCon, community profile based on clustered sequences with UMAPclust and Meshclust; miniCon, community profile based on clustered sequences refined with overlap check; isoCon, community profile based on detected isoforms by IsoCon on the clustered sequences.

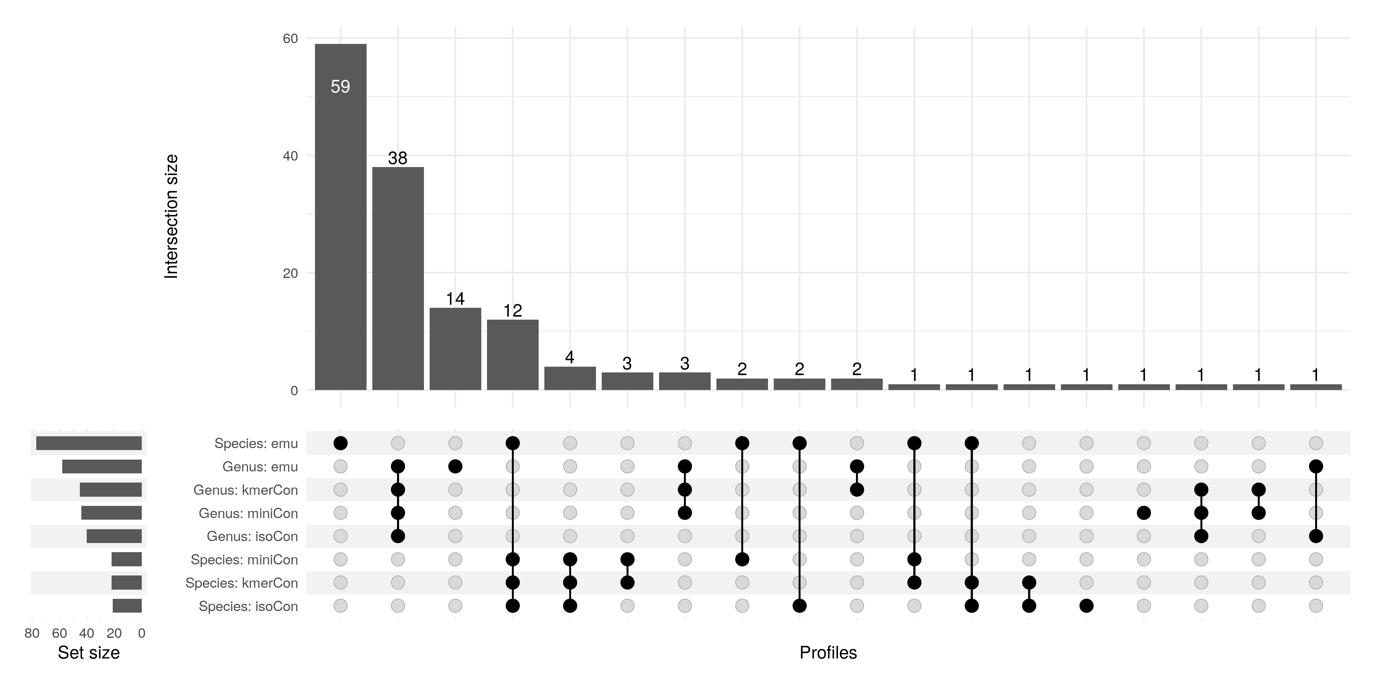

Supplementary Figure 23. Upset plot of the intersection of all identified species and genus tags between various profiles. Emu, read classification profile with Emu; kmerCon, community profile based on clustered sequences with UMAPclust and Meshclust; miniCon, community profile based on clustered sequences refined with overlap check; isoCon, community profile based on detected isoforms by IsoCon on the clustered sequences.

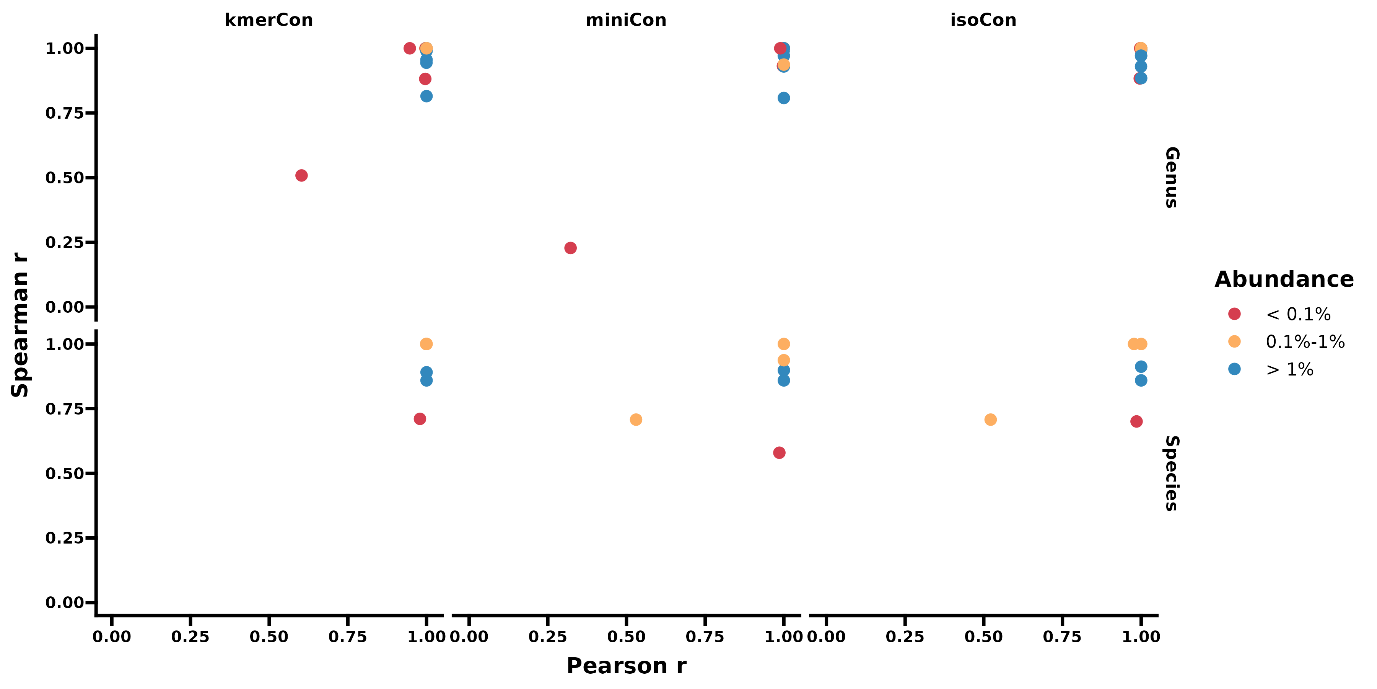

Supplementary Fig. 24: Spearman’s rank and Pearson correlation coefficients of relative abundance of shared genus and species labels (with minimal non-zero values of 30%) between LACA and Emu profile. The points represent the shared taxonomic hits and are colored by the mean abundance in the Emu profile. Emu, read classification profile with Emu; kmerCon, community profile based on clustered sequences with UMAPclust and Meshclust; miniCon, community profile based on clustered sequences refined with overlap check; isoCon, community profile based on detected isoforms by IsoCon on the clustered sequences.

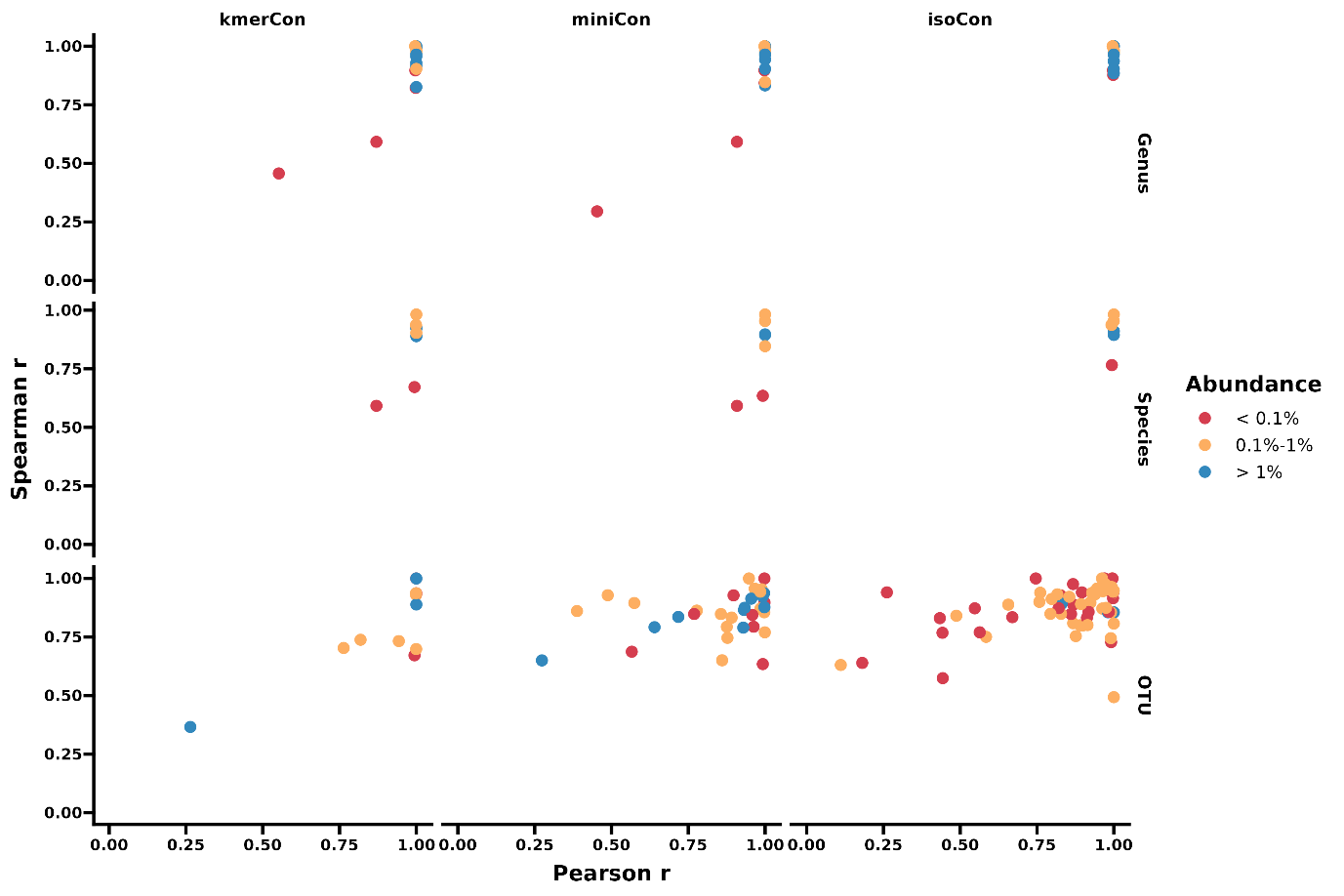

Supplementary Fig.25: Spearman’s rank and Pearson correlation coefficients of relative abundance of shared taxonomic tags (with minimal non-zero values of 30%) between clustering-based and mapping-based abundance. The points represent the shared taxonomic tags and are colored by the mean abundance based on Minimap2 alignments. KmerCon, community profile based on clustered sequences with UMAPclust and Meshclust; miniCon, community profile based on clustered sequences refined with overlap check; isoCon, community profile based on detected isoforms by IsoCon on the clustered sequences.

**Supplementary tables**

Supplementary Table 1. Summarized error profile of corrected sequences with various *de novo* clustering approaches at 200× simulated sequencing coverage

Supplementary Table 2. Quality statistics of ONT Duplex and PacBio CCS consensus sequences generated by isONclust, UMAPclust and Meshclust with a loose identity score of 0.5.

Supplementary Table 3. Quality statistics of consensus sequences generated by UMAPclust and Meshclust with the estimated identity score from the input reads.

Supplementary Table 4. The error profile of various quality-controlled OTU sequences retrieved from the 100,000 subsampled reads from the UMI-tagged amplicon dataset.

Supplementary Table 5. The error profile of various quality-controlled OTU sequences retrieved from the 1,000,000 subsampled reads from the UMI-tagged amplicon dataset.

Supplementary Table 6. Quality statistics of the OTUs from the ZymoBIOMICS 16S rRNA amplicon dataset.

Supplementary Table 7. The cover rate of Emu features in the top BLAST hits of LACA OTU sequences.

Supplementary Table 8. The Emu species and genus features detected in the BLAST hits of LACA OTU sequences.

The min alignment identity and cover of BLAST hits are 0.97 and 0.99, respectively. The mean relative abundance is the mean of non-zero values across all 12 samples.

Supplementary Table 9. The best BLAST hit of the *Peptostreptococcaceae* OTUs against the curated EzBioCloud database for 16S rRNA gene sequences.

Supplementary Table 10. Summary of the uncovered taxonomic features by Emu.

The min alignment identity and cover of BLAST hits are 0.97 and 0.99, respectively. The mean relative abundance is the mean of non-zero values across all 12 samples.
